# c-Myc inhibition is critical for HSC expansion and Rad51 expression

**DOI:** 10.1101/403584

**Authors:** Merve Aksoz, Esra Albayrak, Galip Servet Aslan, Raife Dilek Turan, Lamia Yazgi Alyazici, Pinar Siyah, Emre Can Tuysuz, Serli Canikyan, Dogacan Yucel, Neslihan Meric, Zafer Gulbas, Fikrettin Sahin, Fatih Kocabas

## Abstract

c-Myc plays a major role in the maintenance of glycolytic metabolism and hematopoietic stem cell (HSC) quiescence. Targeting modulators of HSC quiescence and metabolism could lead to HSC cell cycle entry with concomitant expansion. Here we show that c-Myc inhibitor 10074-G5 treatment leads to 2-fold increase in murine LSKCD34^low^ HSC compartment post 7 days. In addition, c-Myc inhibition increases CD34+ and CD133+ human HSC number. c-Myc inhibition leads to downregulation of glycolytic and cyclin-dependent kinase inhibitor (CDKI) gene expression *ex vivo* and *in vivo*. In addition, c-Myc inhibition upregulates major HDR modulator Rad51 expression in hematopoietic cells. Besides, c-Myc inhibition does not alter proliferation kinetics of endothelial cells, fibroblasts or adipose derived mesenchymal stem cells, however; it limits bone marrow derived mesenchymal stem cell proliferation. We further demonstrate that a cocktail of c-Myc inhibitor 10074-G5 along with tauroursodeoxycholic acid (TUDCA) and i-NOS inhibitor L-NIL provides a robust HSC maintenance and expansion *ex vivo* as evident by induction of all stem cell antigens analyzed. Intriguingly, the cocktail of c-Myc inhibitor 10074-G5, TUDCA and L-NIL improves HDR related gene expression. These findings provide tools to improve *ex vivo* HSC maintenance and expansion, autologous HSC transplantation and gene editing through modulation of HSC glycolytic and HDR pathways.

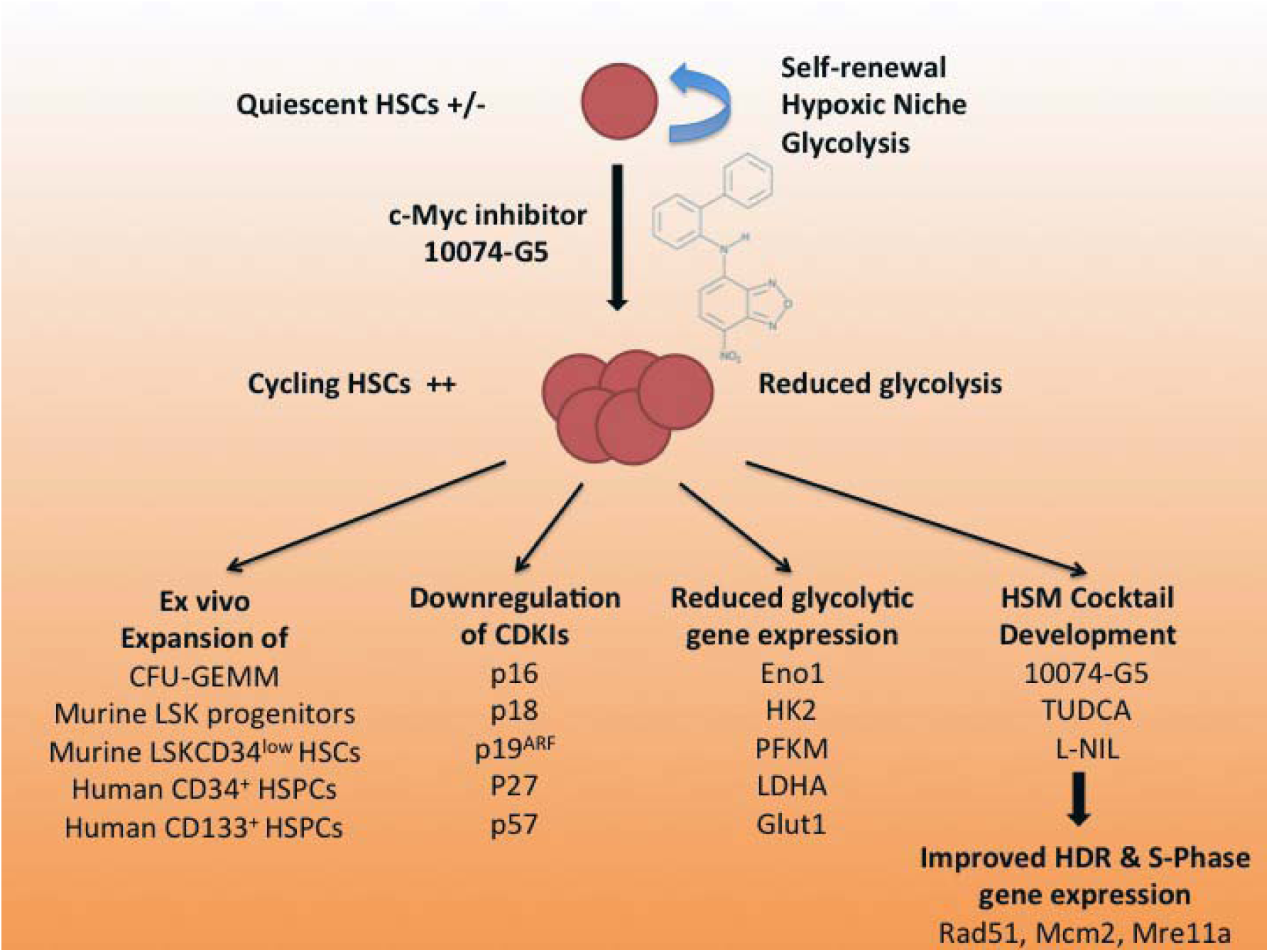
Graphical abstract.

**Highlights:**

- c-Myc inhibition induces ex vivo murine and human hematopoietic stem and progenitor cell proliferation and in vivo murine HSC pool
- c-Myc inhibition downregulates CDKIs and glycolytic gene expression in hematopoietic cells
- c-Myc inhibition do not cause apparent changes to endothelial cells, fibroblasts, or AD-MSCs but it limits BM-MSC proliferation
- c-Myc inhibition along with TUDCA and L-NIL improves HSC maintenance and expansion as evident by induction of all HSC surface antigens analyzed
- c-Myc inhibition alone upregulates major HDR modulator Rad51 expression in hematopoietic cells
- Cocktail of c-Myc inhibitor 10074-G5, TUDCA and L-NIL improves HDR and S-phase related gene expression

## 1. INTRODUCTION

The primarily therapeutic modality for many hematopoietic disorders such as leukemia, lymphoma, some solid cancers and autoimmune disorders is bone marrow transplantation, which relies on the ability of a small number of HSCs. However, lack of HLA-matched donors limits allogeneic HSC transplantation. Even if a donor is available, higher number of HSCs is needed to reduce toxicity of the procedure and for successful engraftment (1). In addition, development of HSC expansion technologies are needed especially post gene editing in HSCs. This issue has been largely addressed by developing *ex vivo* methods to expand HSCs (2, 3).

Earlier studies showed the applicability of *ex vivo* HSC expansion using small molecules (3-8) by targeting cell cycle inhibitors, HSC quiescence regulators and inhibitory factors of *ex vivo* HSC expansion via utilizing hematopoietic small molecules (HSMs). Tauroursodeoxycholic acid (TUDCA), for instance, promotes HSC reconstitution capacity through reducing endoplasmic reticulum stress (9). Besides, it has been reported that highly purified human CD34+ progenitor cells contain iNOS (nitric oxide synthase) and NO showed inhibitory role on human hematopoiesis (10). Treatment with selective iNOS inhibitor, L-N-6-(1-iminoethyl)-lysine hydrochloride (L-NIL), increased preservation of CD34+ progenitors. In addition, α-Tocopherol has been shown to play role in the expansion of primitive hematopoietic cells through its radioprotective property (11).

The *c-Myc* was discovered in human Burkitt’s lymphoma, as a cellular homologue to the viral oncogene (v-Myc) of the avian myelocytomatosis retrovirus (12, 13). C-Myc has been emerged as essential regulator of variety of cellular processes such as proliferation, cell growth, differentiation, angiogenesis and apoptosis (14-16). C-Myc controls energy metabolism by directly activating of genes associated with glycolysis, glutamine metabolism and mitochondrial biogenesis. Intriguingly, Wilson et al. showed that conditional deletion of c-Myc increased lineage negative cells *in vivo.* (17). However, to date, inhibition of c-Myc has not been explored for *ex vivo* or *in vivo* HSC expansion. HSMs become instrumental in developing new approach to expand and maintain functional HSCs *ex vivo* post gene editing studies and may have a potential for improving clinical transplantation of HSCs in the future. We have suggested that by targeting c-Myc inhibitor with 10074-G5 along with several known HSMs, it will be possible to improve maintenance and expansion of HSCs. In this study, c-Myc inhibitor 10074-G5 alone and how it affects the HSC pool, TUDCA, L-NIL and α-Tocopherol, and their combination have been studied in *ex vivo* HSC expansion.

## 2. MATERIALS AND METHODS

### 2.1. Materials

10074-G5 (Calbiochem, Cat no.475957), TUDCA (Calbiochem, Cat no.580549-16M), α-Tocopherol (Sigma Aldrich, Cat no. 258024-5G) and L-NIL (Enzo, Cat no.ALX-270-010-M010) were dissolved in Dimethyl sulfoxide. Dimethyl sulfoxide (% 0.5) used as a control (DMSO, Santa Cruz Biotech, cat. no. sc-3590329). Balb/c (18-20)mice have been used in the studies. All human and animal studies were approved by the Institutional Clinical Studies Ethical Committee of Yeditepe University (Decision numbers 547 and 548), and the Institutional Animal Care and Use Committee of Yeditepe University (YUDHEK, decision number 429).

### 2.2. Bone marrow collection and lineage depletion

Bone marrow cells were harvested from femur and tibia of 6-8 week Balb/c mice (18-20) (YUDETAM, Turkey) following euthanasia. Dissections were performed under the hood. Femur and tibia were cleaned from muscle and connective tissues by scraping with scalpels (7, 21-24). Bone marrow cells were harvested by flushing femurs and tibias with ice cold DPBS using a syringe and a 26G needle. The cell suspension was filtered through 70 µm cell strainer (BD Pharmingen, cat. no. 352350). Magnetic cell separation was performed according to a protocol modified from mouse hematopoietic progenitor (stem) cell enrichment set - DM (BD Pharmingen, cat. no. 558451) (7, 21-24). Cells were resuspended in 0.5 ml ice cold DPBS supplemented with 2% (v/v) fetal bovine serum (FBS, Sigma Aldrich, USA, cat. no. 12103C) and then 5 µl BD Fc Block™ was added and incubated on ice for 15 minutes. 25 µl Biotinylated Mouse Lineage Depletion Cocktail was added to cell suspension incubated on ice for 15 minutes then washed with 10 ml DPBS supplemented with 2% (v/v) FBS and spinned at 1500 rpm for 5 minutes. After removing of supernatant, cells were resuspended in 225 µl DPBS supplemented with 2% (v/v) FBS. Then, cells were labeled with 25 µl BD IMag™ Streptavidin Particles Plus on ice for up to 30 minutes of incubation. Cell suspension were washed with 10 ml DPBS supplemented with 2% (v/v) FBS, spinned at 1500 rpm for 5 minutes and supernatant was removed respectively. Labeled cell suspension is then placed within the magnetic field of the IMagnet (BD Pharmingen cat. no. 552311) by dissolving in 3 ml DPBS supplemented with 2% (v/v) FBS. Labelled cells were transferred to a 12×75 mm round bottom test tube and placed on IMagnet for 10 minutes. After 10 minutes, the supernatant (negative fraction) was aspirated without disturbing the tube and placed in a 2^nd^ new sterile tube for 10 minutes. The positive fraction was resuspended in same volume DPBS supplemented with 2% (v/v) FBS and placed on IMagnet for another 10 minutes. Supernatant was aspirated from 2^nd^ tube and transferred to 15 ml falcon tube after 10 minutes. Supernatant from 1^st^ tube was transferred to new tube and placed on IMagnet for 10 minutes. Supernatant was added to cell suspension in 15 ml falcon tube. The final depleted fraction contains lineage negative (Lin^-^) cells.

### 2.3. Staining of hematopoietic stem cells for flow cytometry analysis

Lineage negative cells (30,000 cells/well) were cultured in a 96 well plate (Corning Costar, cat no. CLS3599) in Serum-Free Expansion Medium (StemSpan SFEM, Stemcell technologies, cat no. 09650) supplemented with SCF (1000 unit/mL), TPO (1000 unit/mL), FLT-3L (5000 unit/mL) (all from R&D Systems Inc., Minneapolis) and 1% (v/v) PSA (10,000 units/ml penicillin and 10,000 ug/ml streptomycin and 25 µg/mL of Amphotericin B, Gibco, cat.no.15240062) and treated with small molecules for 7 days. Following antibodies were used for identification of LSK and LSKCD34^low^ populations: Anti-Mouse CD16/32 Fc block, c-Kit (CD117) PE, CD34 FITC, Sca-1 PE-Cy7 and mouse APC lineage cocktail BD StemFlow, cat no. 560492). Staining of hematopoietic stem and progenitor cells (HSPCs) and flow cytometric analysis were performed as previously described (7, 21-24). Two separate mix was prepared with antibodies given above. As a first step 50 µl mix 1 (5 ml DPBS and 2 µl Fc block) was added to the cells and mixed by pipetting 3-4 times then incubated for 10 minutes at room temperature. 50 µl of mix 2 (5 ml DPBS and 2 µl lineage cocktail, 2 µl Sca-1, 2 µl c-Kit, 2 µl CD34 FITC antibodies) was added and mixed by pipetting 3-4 times then incubated for 15 minutes at room temperature. Cell kinetics was analyzed by flow cytometry (FACSAria III, BD, cat. no. 23-11539-00).

### 2.4. Cell cycle analysis

Murine LSKCD34^low^ (Lin^-^Sca1^+^c-Kit^+^CD34^low^) cells from mouse lineage negative cell population were sorted by flow cytometry (FACSAria III, BD Biosciences). Briefly, murine lineage negative cells were suspended in 500 µl DPBS supplemented with 2% FBS and stained with each 2 µl mouse APC lineage cocktail, anti-mouse c-Kit (CD117) PE, Sca-1 PE-Cy7 and CD34 FITC. Cells were stained on ice for 15 minutes and sorted by flow cytometry. Murine LSKCD34^low^ cells were seeded in the HSC expansion medium at a density of 5,000 cells per well in 96 well-plate. They were treated with the 10074-G5, TUDCA, α-Tocopherol, and L-NIL at 10 µm concentration for murine cells in three replicates. After 3 days of the treatment, the cells were stained with 2 µl of Hoechst 33342 (10 µg/ml) (Sigma Aldrich, USA, cat no. 14533) and incubated in humidified incubator at 37°C and 5% CO_2_ for 30 minutes. Then, 1 µl of Pyronin Y (100 µg/ml) (Sigma Aldrich, USA, cat no. P9172-1G) was added and incubated in humidified incubator at 37°C and 5% CO_2_. Stained cells were transferred to sterile flow tubes and cell cycle kinetics were analyzed by flow cytometry as we have done previously (7, 21-24).

### 2.5. Apoptosis analysis

Murine LSKCD34^low^ cells were isolated by flow cytometry from lineage negative cells (FACSAria III, BD Pharmingen) as we have done previously (7, 21-24). Murine LSKCD34^low^ cells were seeded at a density of 5000 cells/well in 96-well plates and treated with HSMs in three replicates at a final concentration of 10 µm for 3 days in the humidified incubator at 37°C and 5% CO_2._ After 3 days, cells were collected from 96-well plates and centrifuged at 1500 rpm for 5 minutes. After removing of supernatant, cells were suspended in 50 µl 1X binding buffer (BD Pharmingen, cat no 556570) and then stained according to the manufacturer’s manual (BD Pharmingen, cat no 556570) with 1 µl FITC Annexin V and 1 µl PI at room temperature for 15 minutes. 200 µl 1X binding buffer were added onto cells and mixed by pipetting. Samples were analyzed by flow cytometry (FACS Aria III, BD Biosciences).

### 2.6. Colony forming unit (CFU) assays

Murine lineage negative (Lin^-^) cells (30.000 cells/well) were treated with 10074-G5, TUDCA, α-Tocopherol, and L-NIL at 10 µm concentration as we have done previously (7, 21-24). After 7 days of treatment, the cells were harvested and counted. Equal numbers of cells were plated in methylcellulose-containing medium (MethoCult™ GF M3434, Stemcell Technologies, Cat.No. 03444) at a density of 65,000 cells per well in 6-well plate, performed in triplicate. After 13 days, CFU-GEMM, CFU-G/M, BFU-E colonies were analyzed by inverted microscope.

### 2.7. Isolation of mononuclear cells from umbilical cord blood (UCB) and flow cytometry analysis of HSC markers

UCB mononuclear cells were isolated by Ficoll-Paque (Histopaque™, Sigma, cat. no.10831) density gradient centrifugation as we have done previously (7, 21-24). Briefly, 15 ml cord blood was diluted with DPBS as 1:1 proportion in a 50 ml falcon tube and mixed gently by inverting. Cord blood was underlaid with 15 ml Ficoll-Paque and centrifuged at 3000 rpm for 15 minutes without brake. After centrifugation upper phase was removed and cloudy interphase transferred to new 50 ml falcon tube. Mononuclear cells were washed with 3x volume of DPBS and mixed by gentle inverting. Cells were centrifuged at 1500 rpm for 5 minutes with brake, and then supernatant was removed. Cell pellet was suspended in 10 ml DPBS and cell count was assessed by counting on hemocytometer. UCB mononuclear cells were plated in 96 well-plate in human HSC expansion medium at 10,000 cells per well for flow cytometry analysis (25). The expansion medium consists of Serum-Free Expansion Medium (StemSpan™ Serum-Free Expansion Medium (SFEM), Stemcell Technologies, cat. no. 09650) supplemented with human cytokine cocktail (StemSpan™ CC100, Stemcell Technologies, cat. no. 02690) and 1% PSA (10,000 units/ml penicillin and 10,000 µg/ml streptomycin and 25 µg/mL of Amphotericin B, Gibco, cat. no.15240062). After 7 days of small molecules treatments (1 µM), the UCB mononuclear cells were labelled with PE-conjugated anti-human CD34 (Biolegend, Cat.No.343506), and APC-conjugated anti-human CD133 (Miltenyibiotec, Order No.130-090-826) antibodies according to the manufacturer’s manual (Stemcell Technologies). CD34 and CD133 human HSC surface markers were analyzed by flow cytometry (FACSAria III, BD, cat. no. 23-11539-00) as we have done previously (25).

### 2.8. Isolation of adipose derived mesenchymal stem cells (AD-MSC)

60 ml adipose tissue from liposuction sample was placed into a 500 ml bottle with same amount of collagenase solution. The tissue was digested at 37°C for 1 hour by continuous shaking. The digested tissue was centrifuged at 2500 rpm for 7 min at room temperature (RT). The adult adipocytes and collagenase in the supernatant were discarded. The remaining pellet was resuspended with 2 ml of erythrocyte lysis buffer. The cell suspension was completed to 50 ml with erythrocyte lysis buffer and incubated at 37°C for 10 min by continuous shaking. The cell suspension was centrifuged at 1400 rpm for 7 min at RT and supernatant was discarded. The pelleted cells were washed with 1X PBS then centrifuged at 1400 rpm for 7 min at RT, and supernatant were discarded. The cells were resuspended in 6-8 ml DMEM (Gibco) then filtered through 100 µm cell strainer. The cells seeded onto tissue culture polystyrene flasks at the density of 10^6^cells/150 cm^2^.

### 2.9. Isolation of bone marrow derived mesenchymal stem cells (BM-MSC)

Mouse bone marrow mesenchymal stem cells were isolated according to a protocol modified from Soleimani and Nadri (26). Bone marrow cells were collected as described above and seeded at a density of 30 × 10^6^ cells in in T75 cm^2^ flasks (Sigma Aldrich, cat no. CLS3290) in Dulbecco’s Modified Eagle’s Medium (DMEM, Gibco) supplemented with 15% (v/v) FBS (Sigma Aldrich, USA, cat no. 12103C) and 1% (v/v) PSA (10,000 units/ml penicillin and 10,000 µg/ml streptomycin and 25 µg/mL of amphotericin B, Gibco, Cat.No.15240062). One day after, non-adherent cells were removed and medium was changed in every 3-4 days. After 15 days of initiating culture, MSCs were lifted with 0.25% tyripsin-EDTA (Sigma Aldrich, USA, cat no. 25200056) and cultured in T75 cm^2^ flasks. 10,000 BM-MSCs per well were seeded on a 96-well plate for small molecule treatments.

### 2.10. Analysis of cell proliferation

Human dermal fibroblast (HDF) cells (kindly provided by Prof. Dr. Dilek Telci from Yeditepe University, Turkey) were seeded at a density of 5000 cells per well in 96-well plates. Human umbilical vein endothelial cells (HUVECs, ATCC^®^ CRL1730™) were seeded on 96 well plates at a density of 2,000 cells per well. BM-MSCs and AD-MSCs were seeded in 96 well plates at a density of 5,000 cell and 10,000 cell per well respectively. Next day, cells were treated with HSMs for three days. Three days after, cell medium was removed. WST1 reagent were diluted in 1:10 range with culture medium (Cell Proliferation Reagent WST-1, Roche, cat. no. 11644807001) and added as 50 µl for each sample. Plates were incubated in humidified incubator at 37°C and 5% CO_2_ in dark and measured the absorbance of the samples at the time point of 1 hour, 2 hours and 3 hours using a microplate reader (Thermo Scientific) at 420-480 nm as we have done previously (27).

### 2.11. Analysis of HSC content in the bone marrow and peripheral blood

For *in vivo* analysis of HSC proliferation and gene expression, 10074-G5 was diluted in 100 µl DPBS at a concentration of 1 µM and injected into adult wild type Balb/c mice intraperitoneally on 1^st^, 4^th^ and 7^th^ days. At 10^th^ day, mice were euthanized and bone marrow cells were harvested from femur and tibia. Bone marrow cells were analyzed by flow cytometry following staining with APC lineage cocktail, c-Kit (CD117) PE, Sca-1 PE-Cy7, CD150 FITC and CD48 APC. Moreover, to measure HSC mobilization peripheral blood was collected through retroorbital bleeding and stained with APC lineage cocktail, c-Kit (CD117) PE, Sca-1 PE-Cy7, CD34 FITC then analyzed by flow cytometry (BD Calibur) as we have done previously (7, 21-24).

### 2.12. Gene expression analysis

Gene expression analysis was carried as we have done previously in hematopoietic compartments (7, 21-24). Briefly, whole bone marrow (WBM) was harvested from Balb/c mouse as described before. LSKCD34^low^ cells were isolated by FACS Aria III (BD Biosciences). Supernatant was removed and cell pellet was stored at −80°C. WBM cells were separated for further RNA isolation and analysis of c-Myc expression by real-time PCR. Additionally, 2×10^6^ Lin^-^ cells were seeded on a 6-well plate and treated with DMSO control (%0.5) and 10 µM dose of 10074-G5 for 5 days in 37°C humidified incubator. Following treatment, the cells were collected for RNA isolation in 15 ml falcon tubes at −80°C. RNA was isolated according to the manufacturer’s protocol GenElute™ Mammalian Total RNA Miniprep Kit (Sigma Aldrich, Cat no. RTN70).

RNAs were converted to cDNA by using ProtoScript^®^ First Strand cDNA Synthesis Kit (NEB, Cat no. E6300S). All eluted RNA was used as a template and upto 1 µg cDNA was synthesized. Each RNA sample was mixed with 2 µl random primer and denatured at 65°C for 5 minutes and then transferred into ice. Predesigned primers (**Table 1**) obtained from NIH Mouse depot (https://mouseprimerdepot.nci.nih.gov/) ordered from Sentegen Biotechnology, Turkey. Maxima SYBR Green qPCR Master Mix (2X) (Thermo Scientific, Cat. No. K0222) was used in order to perform real-time PCR. Reaction was conducted on BioRad CFX96 Touch™ real-time PCR Detection System. β-actin was used as an internal control. Data were analyzed by using 2-ΔΔCt Method.

**Table 1.**
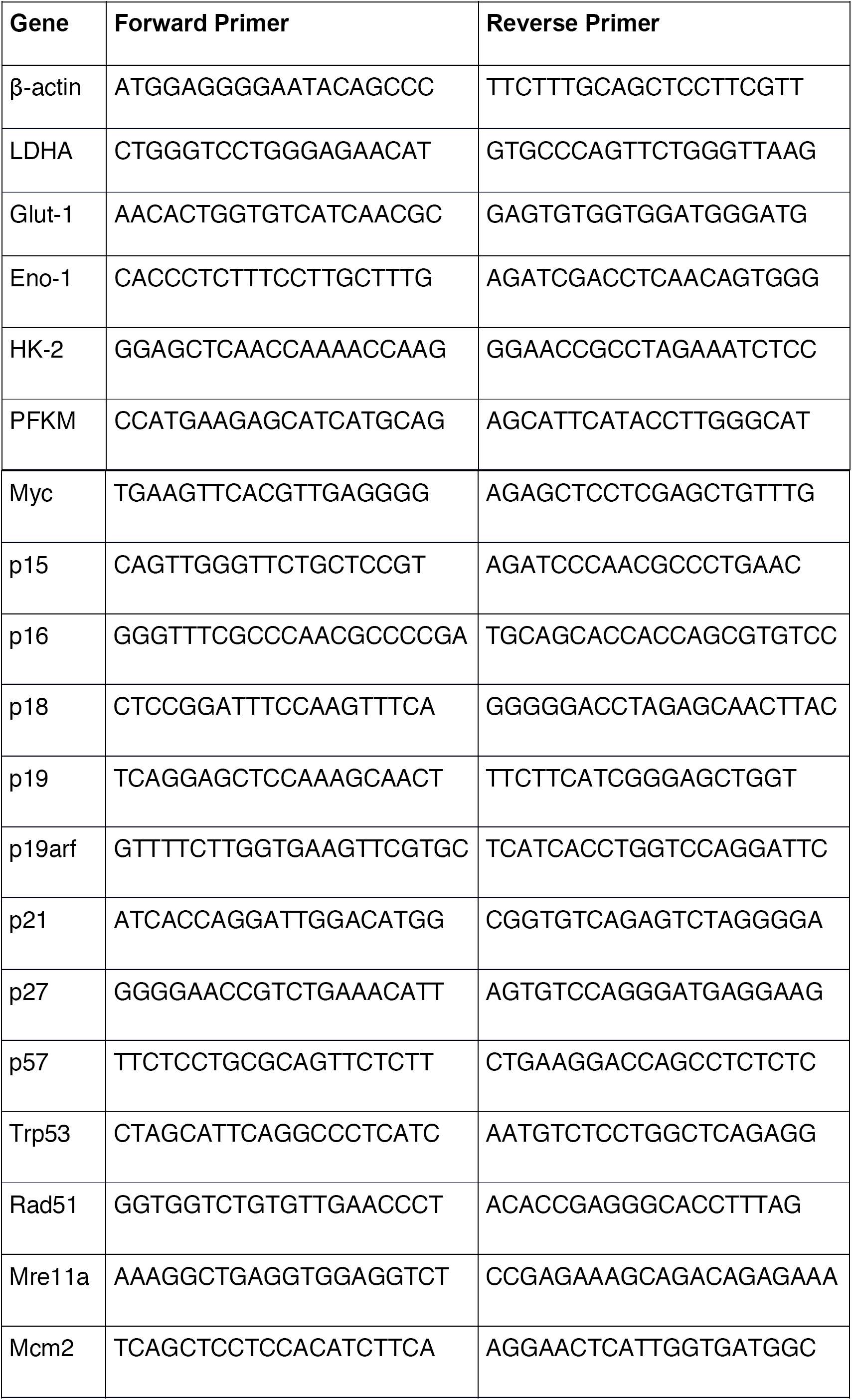
List of primers used for real-time PCR

### 2.14 Combinatorial hematopoietic small molecule (HSM) treatments

Small molecule master mixtures were prepared as shown in the **Table 2**. 30,000 lineage negative cells per well were treated with small molecule mixtures in 96-well plates at a final concentration of 10 µM, and HSC content was analyzed by flow cytometry after 7 days.

**Table 2.** Mixtures of hematopoietic small molecules

|  | M1 | M2 | M3 | M4 | M5 | M6 | M7 | M8 | M9 | M10 | M11 |
| --- | --- | --- | --- | --- | --- | --- | --- | --- | --- | --- | --- |
| 10074-G5 | + | + | + | - | - | - | + | + | + | - | + |
| TUDCA | + | - | - | + | + | - | + | + | - | + | + |
| α-Tocopherol | - | + | - | + | - | + | + | - | + | + | + |
| L-NIL | - | - | + | - | + | + | - | + | + | + | + |

### 2.15 Repopulation Analysis

Murine HSCs (LSKCD34^low^) have been isolated by FACS (FACSAria III, BD Biosciences) and treated with DMSO, 10074-G5, and Mixture M8 in HSC expansion medium for 7 days. About 500 ex vivo expanded HSCs have been transplanted into NOD/SCID mice. Engraftment and repopulation of transplanted HSCs have been studied at different time points. Flow cytometry analysis has been performed to confirm multilineage reconstitution. For analysis of repopulation of mouse HSCs, peripheral blood cells of recipient NOD/SCID CD45.1 mice have been collected by retro-orbital bleeding, followed by lysis of red blood cells and staining with anti-CD45.2-FITC, anti-CD45.1-PE, anti-Thy1.2-PE (for T-lymphoid lineage), anti-B220-PE (for B-lymphoid lineage), anti-Mac-1-PE, or anti-Gr-1-PE (cells co-staining with anti-Mac-1 and anti-Gr-1 are deemed to be of the myeloid lineage) monoclonal antibodies (BD Pharmingen). The “percent repopulation” have been determined based on the staining results of anti-CD45.2-FITC and anti-CD45.1-PE as we have done previously (7, 21-24).

### 2.16 Statistical Analysis

Results are expressed as mean ± SEM and a 2-tailed Student *t* test was used to determine the level of significance. p < 0.05 was considered statistically different.

## 3. RESULTS

### 3.1. *Ex vivo* c-Myc inhibition induces mouse HSC maintenance and expansion

We treated Lin^-^ cells *ex vivo* with c-Myc inhibitor and analyzed the frequency of HSCs following 7 days of expansion procedure by flow cytometry after staining with corresponding surface antigens. C-Myc inhibitor 10074-G5 significantly increased LSK progenitor and LSKCD34^low^ HSC compartment compared to DMSO control (p < 0.01) **(Figure 1)**. Functional expansion of HSCs following c-Myc inhibitor 10074-G5 has been studied with colony forming assays. To this end, Lin^-^ cells have been treated with DMSO (control, 0.5%), c-Myc inhibitor 10074-G5 (10 µM) for 7 days in culture, then methylocellulose colony forming assays were performed. We demonstrated that c-Myc inhibitor 10074-G5 treated lineage negative cells had higher number of primitive progenitor and hematopoietic stem cells as evident by increased number of CFU-GEMM colonies and erythroid progenitor cells (BFU-E colonies) compared to DMSO control but no significant change was observed in the number of myeloid progenitor cells (CFU-G/M/GM colonies) (**Figure 2**).

**Figure 1.**
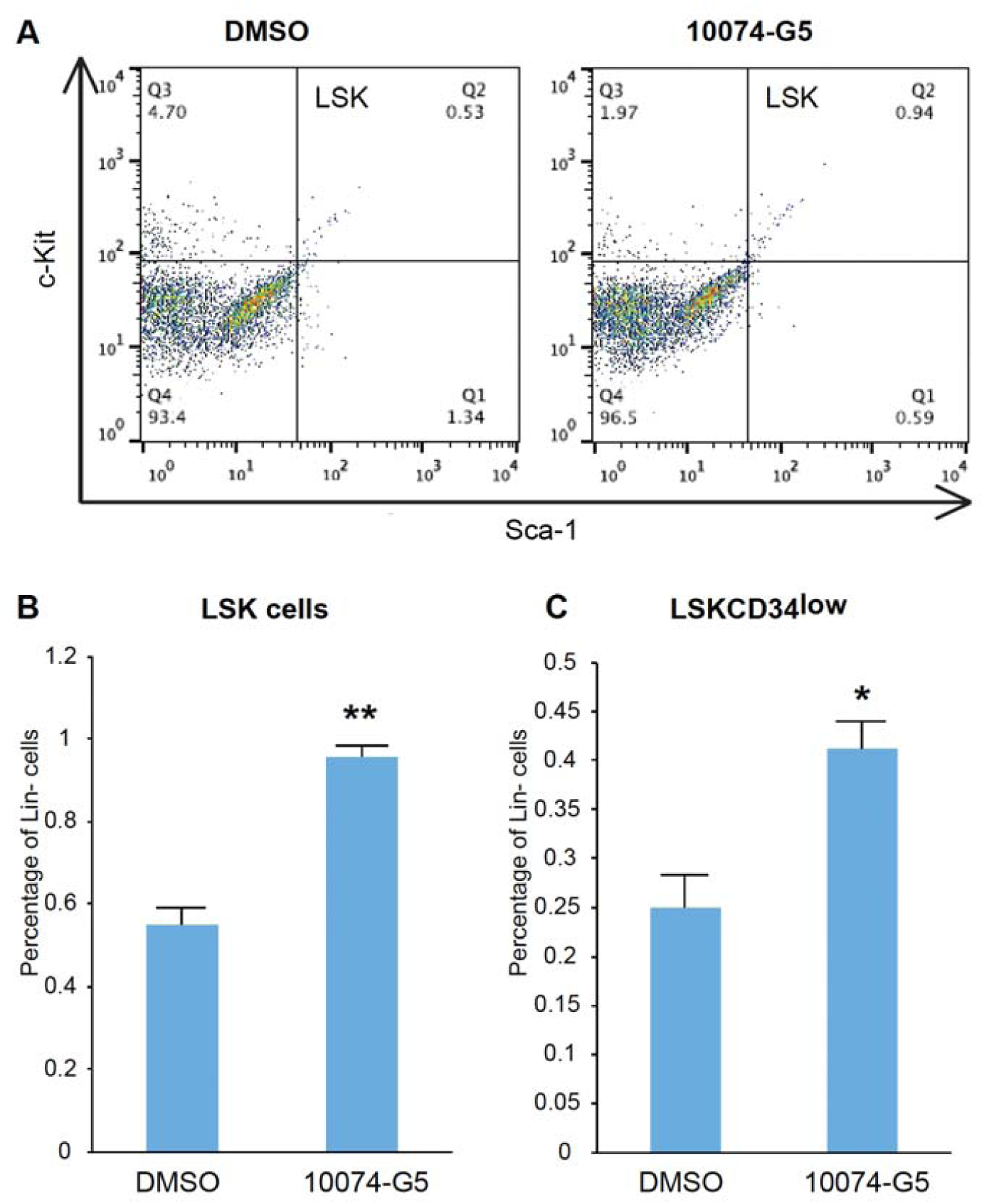
Effect of c-Myc inhibition in murine HSC content. Mouse Lin-cells have been treated with DMSO (control, 0.5%) and c-Myc inhibitor 10074-G5 (10 µM) for 7 days, performed **A)** flow cytometry and determined percent of **B)** Lin^-^Sca1^+^c-Kit^+^ (LSK) and **C)** Lin^-^Sca1^+^c-Kit^+^CD34^low^ (LSKCD34^low^) cells. * p < 0.05, ** p<0.01. n=3.

**Figure 2.**
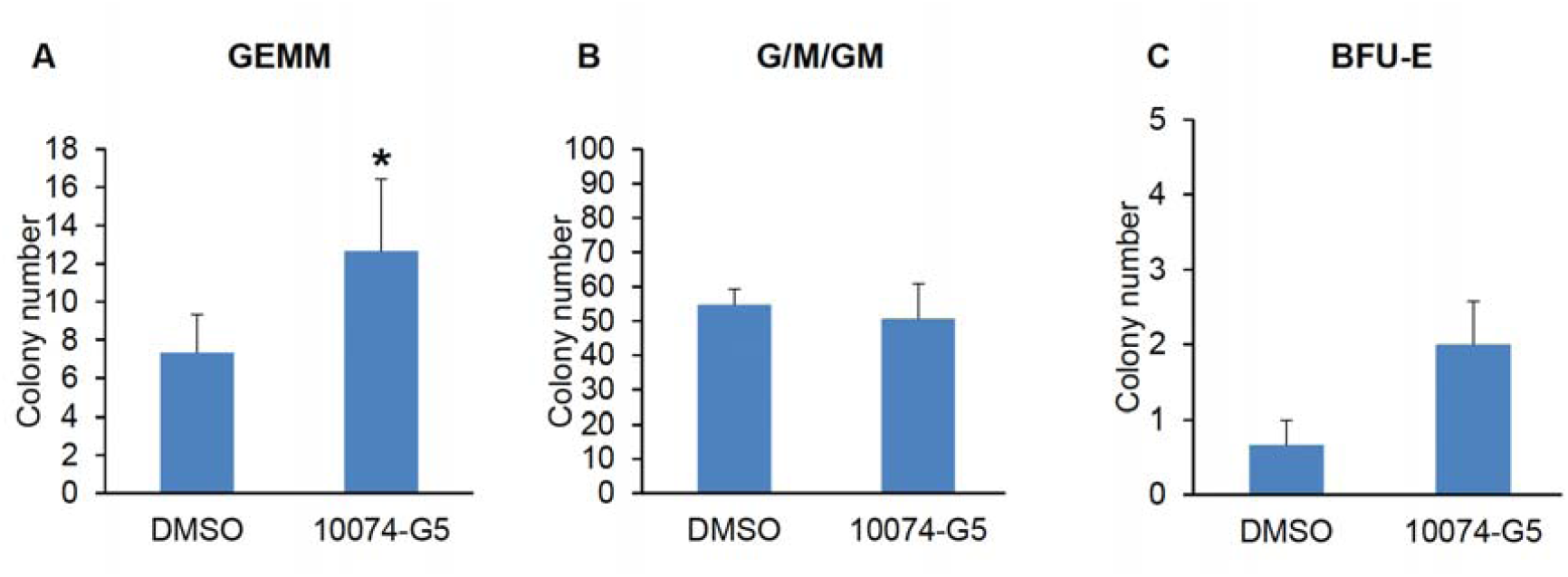
Colony forming assay. Murine Lin-cells were treated with DMSO and c-Myc inhibitor 10074-G5 for 7 days and analyzed the number of **A)** GEMM, **B)** G/M/GM, and **C)** BFU-E colonies formed post 13 days. * p < 0.05. n=3

HSC expansion following c-Myc inhibition was further confirmed by cell cycle analysis. We examined the cell cycle of LSKCD34^low^ HSCs by using Hoechst 33342 and Pyronin Y staining. We found a slight increase of HSC content in G_1_ phase of the cell cycle when treated with 10074-G5 (**Figure S1A**). This indicated that 10074-G5 treated HSCs were much less quiescent and prone to proliferation. Next, we examined apoptosis status of HSCs by Annexin V and propodium iodide (PI) staining. We did not detect any significant apoptotic cells in 10074-G5 treated HSCs compared to control (**Figure S1B**).

### 3.2. Analysis of gene expression profile post treatment with 10074-G5

To understand how inhibition of c-Myc contributes to expansion of hematopoietic stem and progenitor cell, we analyzed expression of c-Myc, c-Myc target genes and cyclin-dependent kinase inhibitors (CDKI) following *ex vivo* treatments of Lin^-^ cells with 10074-G5. Thus, Lin^-^ cells were treated with 10074-G5 at a concentration of 10 µM and DMSO control (%0.5) for 5 days in humidified incubator at 37°C and 5% CO_2._ Then, transcription of c-Myc gene, CDKIs and c-Myc targets were analyzed by using real-time PCR. We found that expression of c-Myc was decreased when it was treated with 10074-G5 in comparison to DMSO control. Inhibition of c-Myc activity in Lin^-^ cells *ex vivo* with 10074-G5 treatment was evident by downregulation of c-Myc target genes such as LDHA, Eno1, and Glut1 (**Figure 3A**). In addition, we found that expression level of cell cycle inhibitors including p18 (CDKN2C), p19 (CDKN2D), p21^CIP1^ (CDKN1A), p57^KIP1^ (CDKN1C) were declined (**Figure 3B**).

**Figure 3.**
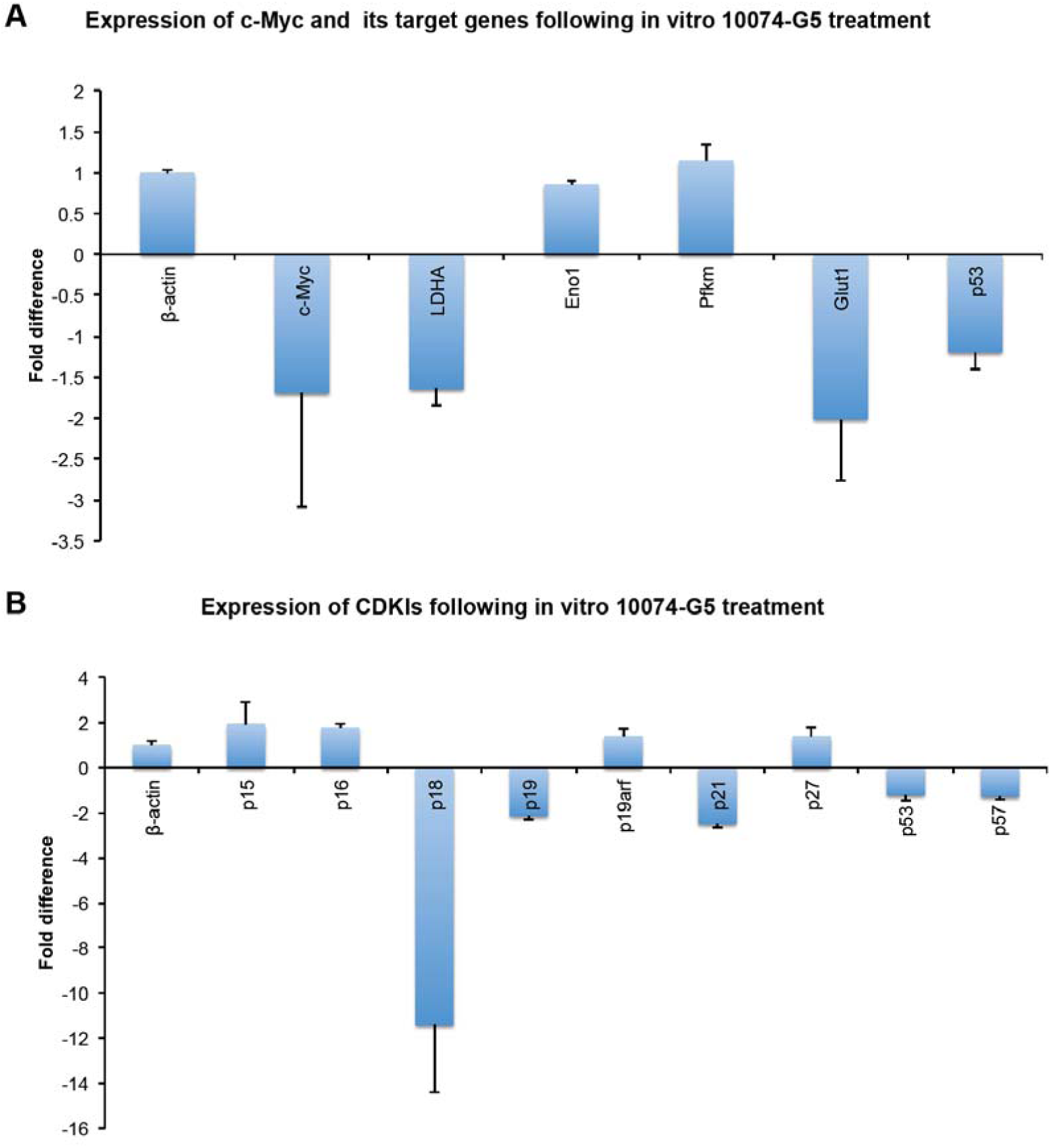
Analysis of gene expression profile post in vitro c-Myc inhibitor 10074-G5 treatments. Expression analysis of A) c-Myc and c-Myc target genes, B) cyclin dependent kinase inhibitors (CDKI) post c-Myc inhibitor 10074-G5 treatment. n=3.

### 3.3. *In vivo* c-Myc inhibition increases HSC content in the bone marrow

Direct *in vivo* targeting of HSCs could provide a physiological approach to modulate HSC activity as an alternative to HSC isolation and *ex vivo* expansion. *In vivo* injection of small molecules could play a molecular switch for expansion and mobilization of HSCs by inhibiting HSC quiescence regulators. To address this, c-Myc inhibitor 10074-G5 was injected to adult wild type mice intraperitoneally. After three doses of injection, stem cell expansion in bone marrow and stem cell mobilization to peripheral blood were examined by flow cytometry (**Figure 4A**). We determined more than 2 fold increase in HSCs content (LSKCD48^-^CD150^+^ cells) in the bone marrow (**Figure 4B**). In addition, we analyzed the extent of any HSC mobilization to peripheral blood by quantification of HSPCs (LSK cells) by flow cytometry (**Figure 4C**). Analysis of peripheral blood HSC content by flow cytometry revealed that there is no significant change in HSC content in the peripheral blood.

**Figure 4.**
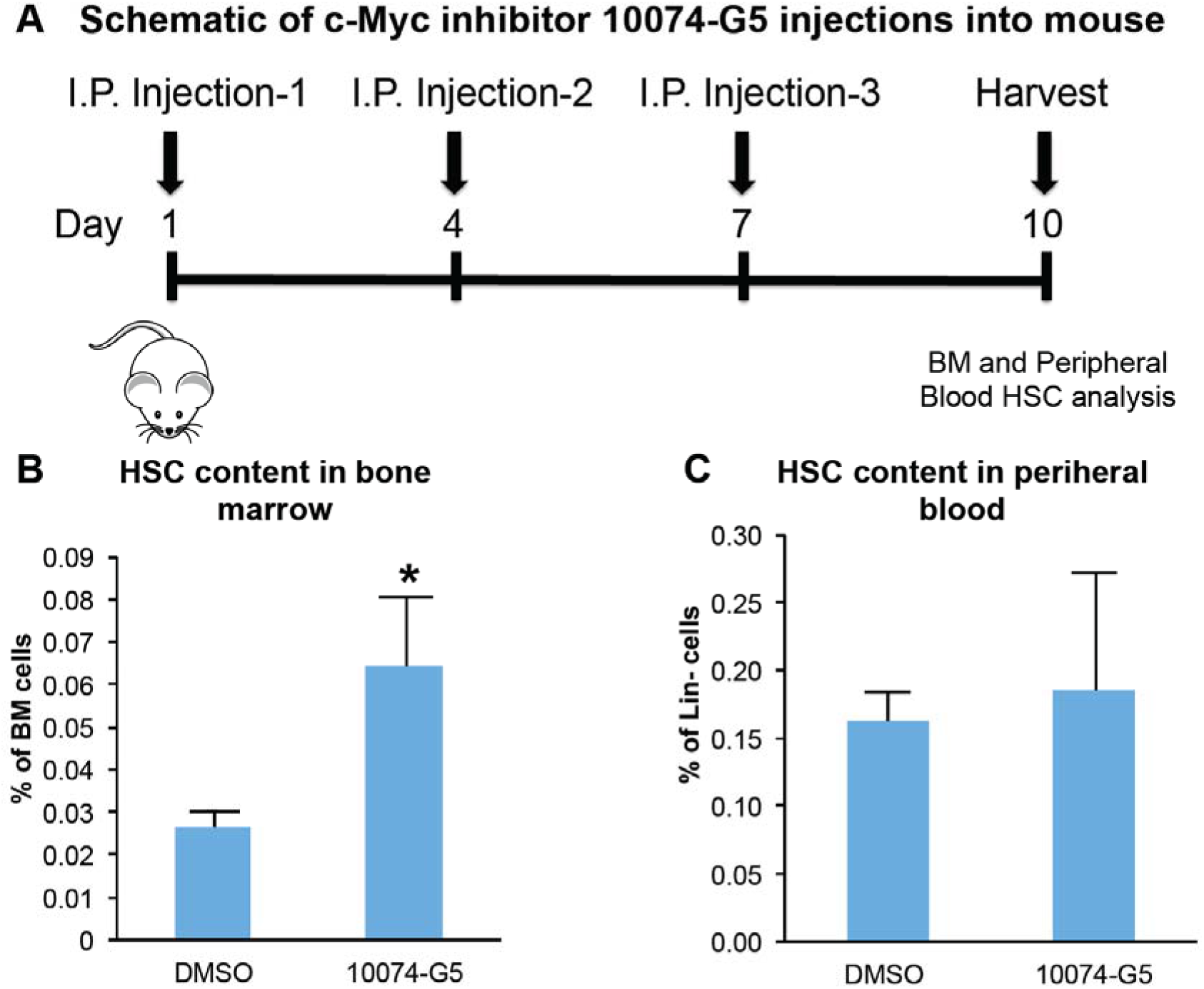
Effect of *in vivo* c-Myc inhibition in the frequency of HSC compartments in the bone marrow and peripheral blood. **A)** Schematic of c-Myc inhibitor 10074-G5 injections into mouse. Percent of B) LSKCD48^-^CD150^+^ HSCs in the bone marrow and C) LSKCD34^low^ HSCs in the peripheral blood. * p < 0.05. n=3.

### 3.4. *In vivo* effect of c-Myc inhibition in expression profile of c-Myc target genes in glycolysis and CDKIs

We, first, confirmed c-Myc gene expression in HSCs (LSKCD34^low^) and compared to the whole bone marrow (WBM) mononuclear cells (**Figure S2**). Next, we analyzed the expression profile of selected c-Myc target genes and cell cycle inhibitors in bone marrow by using RT-PCR following *in vivo* injection of c-Myc inhibitor 10074-G5. We confirmed that c-Myc target genes such as Eno1, HK2, Pfkm and Glut1 were downregulated significantly post c-Myc inhibition (**Figure 5A**). In addition, analysis of cyclin dependent kinase inhibitors (CDKIs) showed the downregulation of p15^INK^4^B^ (CDKN2B), p16^INK^4^A^, p19^ARF^, p21^CIP1^ (CDKN1A), p27^KIP1^ (CDKN1B) and p57^KIP2^(CDKN1C) expression post c-Myc inhibition (**Figure 5B**). These findings support the notion that HSC exit from G0 phase of the cell cycle and proliferate *in vivo*. In addition, downregulation of c-Myc target genes Eno1, HK2, Pfkm and Glut1 indicates a metabolic shift from cytoplasmic glycolysis to mitochondrial phosphorylation. This further suggests that HSCs are becoming more metabolically active state when they proliferate. Similar phenomenon was prevalent in Meis1 or Hif-1α loss of function studies, where we have shown the metabolic shift from cytoplasmic glycolysis to mitochondrial phosphorylation and concomitant increase in HSC pool (7, 21, 22).

**Figure 5.**
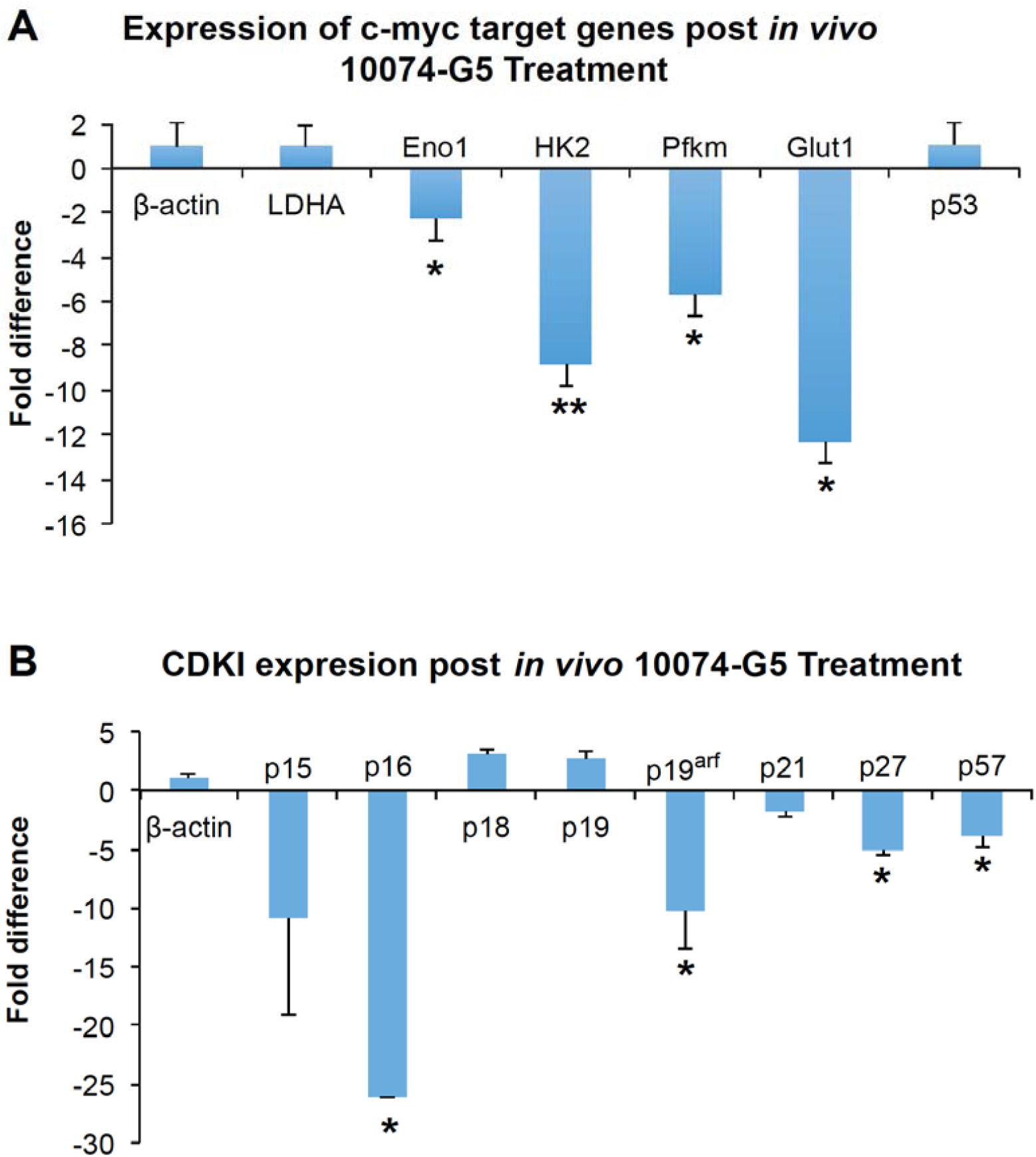
Effect of in vivo c-Myc inhibition in gene expression. After three dose *of* injection of 10074-G5 into mice, bone marrow cells were analyzed for expression of **A)** c-Myc target genes and **B)** CDKIs. Note that *in vivo* injection of c-Myc inhibitor 10074-G5 downregulates the expression of a number of CDKIs and genes that are downstream of c-Myc. * p < 0.05, ** p<0.01. n=3.

### 3.5. *Ex vivo* c-Myc inhibition induces human HSPC maintenance and expansion

To determine the effect of c-Myc inhibitor 10074-G5 in the human HSPC expansion, we treated human umbilical cord blood mononucleated cells with DMSO (0.5%, control) and 1 µM of c-Myc inhibitor 10074-G5 for 7 days. We found that c-Myc inhibition robustly increased human CD34+ and CD133+ HSPC ratio and cell count compared to control as analyzed by flow cytometry (**Figure 6A-D**). In addition, UCB mononuclear cell count increased after c-Myc inhibitor 10074-G5 treatment. Similarly, we have achieved induction of human HSC pool in mobilized peripheral blood cells (**Figure S3**)

**Figure 6.**
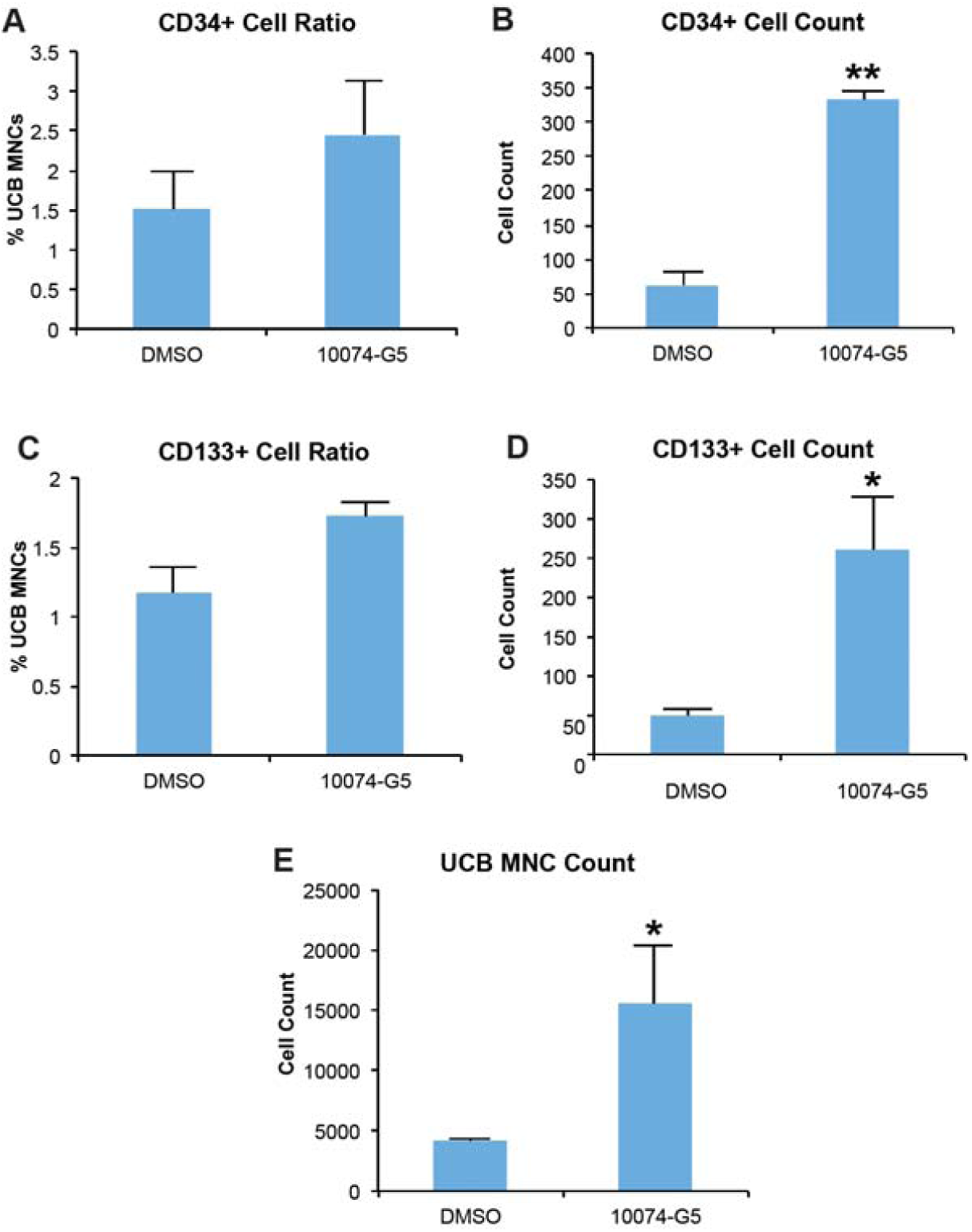
Effect of c-Myc inhibition in human HSPCs. Human UCB mononuclear cells (MNCs) have been treated with DMSO (control, 0.5%) and c-Myc inhibitor 10074-G5 for 7 days and determined A) CD34+ cell ratio, B) CD34+ cell count, C) CD133+ cell ratio and D) CD133+ cell count in UCB MNCs, and E) total UCB MNC count. * p < 0.05, ** p<0.01. n=3.

### 3.6. C-Myc inhibition limits BM-MSC proliferation but do not change proliferation kinetics of AD-MSCs, endothelial cells or fibroblasts

Next, we sought to determine whether the effect of c-Myc inhibitor 10074-G5 is preferentially specific to HSC expansion. Thus, we studied the effect of c-Myc inhibitor 10074-G5 in fibroblast, endothelial cell, adipose derived MSC (AD-MSC), and bone marrow derived MSC (BM-MSC) proliferation (**Figure 7**). Cell proliferation was determined using the WST1 assay after three days of 10074-G5 treatments. We treated dermal fibroblasts (HDF) and human vascular endothelial cells (HUVEC) with c-Myc inhibitor 10074-G5. 10074-G5 treated HUVECs and HDFs showed no difference in proliferation compared to control. In addition, we did not detect any significant change in proliferation kinetics of AD-MSCs compared to control. On the other hand, c-Myc inhibitor 10074-G5 decreased the proliferation rate of BM-MSCs. These results suggest that c-Myc inhibitor do not change proliferation kinetics of AD-MSCs but intriguingly they may suppress the proliferation of BM-MSCs. Taken together, these results indicated that c-Myc inhibitor, 10074-G5, is specific to induction of hematopoietic stem and progenitor cells.

**Figure 7.**
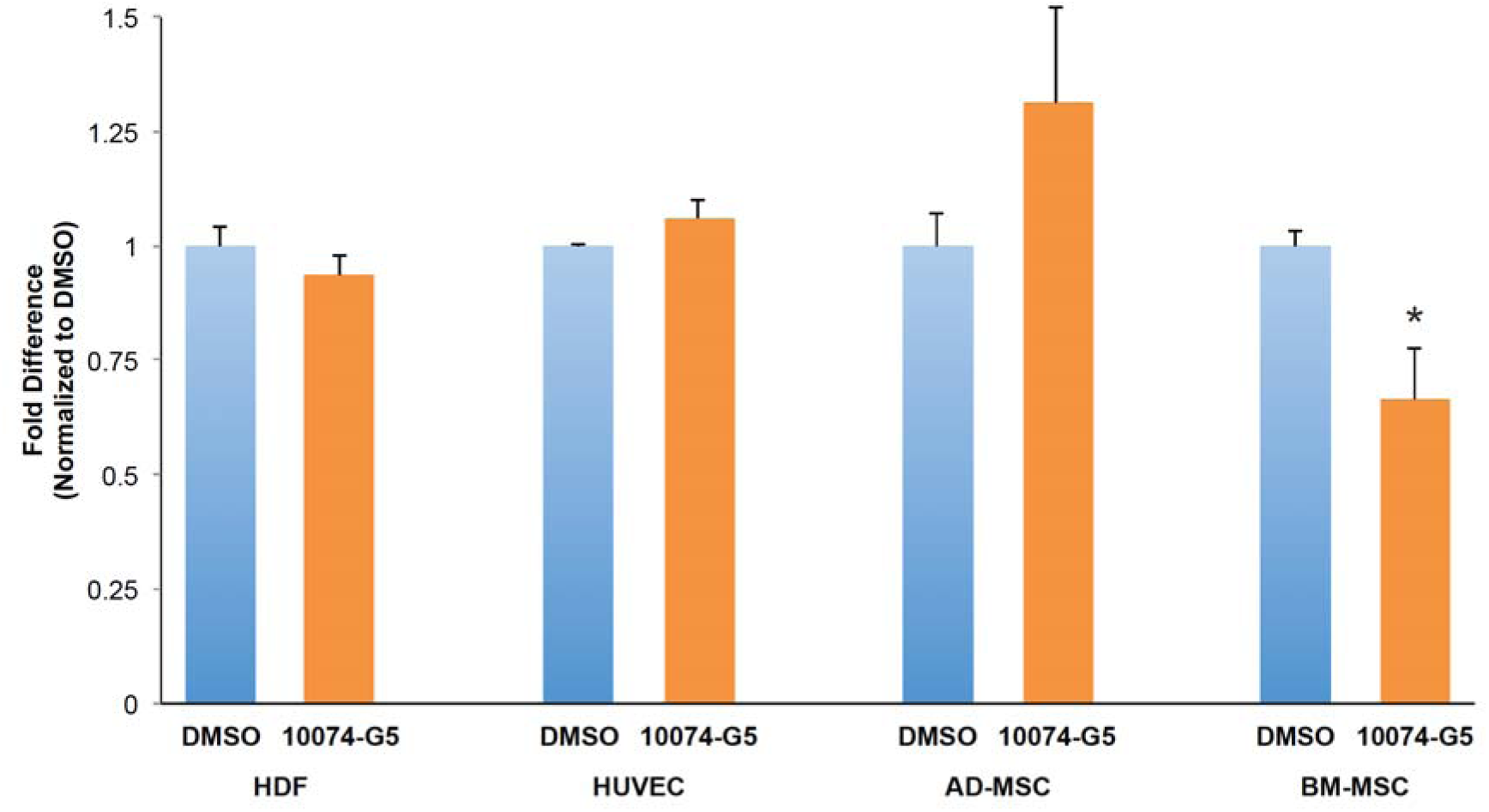
Effect of c-Myc inhibitor 10074-G5 in the proliferation kinetics of fibroblasts, endothelial cells, AD-MSCs and BM-MSCs. HDFs, HUVECs, AD-MSCs and BM-MSCs were treated with DMSO (control, 0.5%) and c-Myc inhibitor 10074-G5 for 3 days and examined by WST1 assay. Fold difference in WST1 absorbance at 450 nm was determined. * p < 0.05. n=3.

### 3.7. Analysis of HSMs in HSPC maintenance and expansion

We hypothesized that an optimum combination of hematopoietic small molecules (HSMs) could provide a robust HSC expansion by targeting several pathways. To this end, we studied previously reported HSMs namely TUDCA, α-Tocopherol, and L-NIL. We show that TUDCA, α-Tocopherol and to some extent L-NIL led to increased frequencies of LSK and LSKCD34^low^ HSC compartment (**Figure 8**). Tested HSMs did not induce c-Kit+ hematopoietic cell content alone (**Figure S4A**) but TUDCA, α-Tocopherol, and L-NIL increased Sca1+ expression (**Figure S4B**). Murine HSC expansion following selected HSM treatments were further confirmed by cell cycle analysis. We found that a decreased HSC content in G_0_ phase and an increased HSC content in G_1_ phase of the cell cycle (**Figure S5**) when treated with α-Tocopherol and L-NIL. Besides, apoptosis analysis of HSM treated HSCs showed that TUDCA, α-Tocopherol and L-NIL do not lead to any significant apoptosis cells (**Figure S6**). In addition, we found that that TUDCA and L-NIL treated lineage negative cells formed higher number of mixed colonies (CFU-GEMM), post 13 days of treatment (**Figure S7A**). Number of CFU-G/M/GM and BFU-E colonies were higher in TUDCA and α-Tocopherol treated hematopoietic cells (**Figure S7B and S7C**).

**Figure 8.**
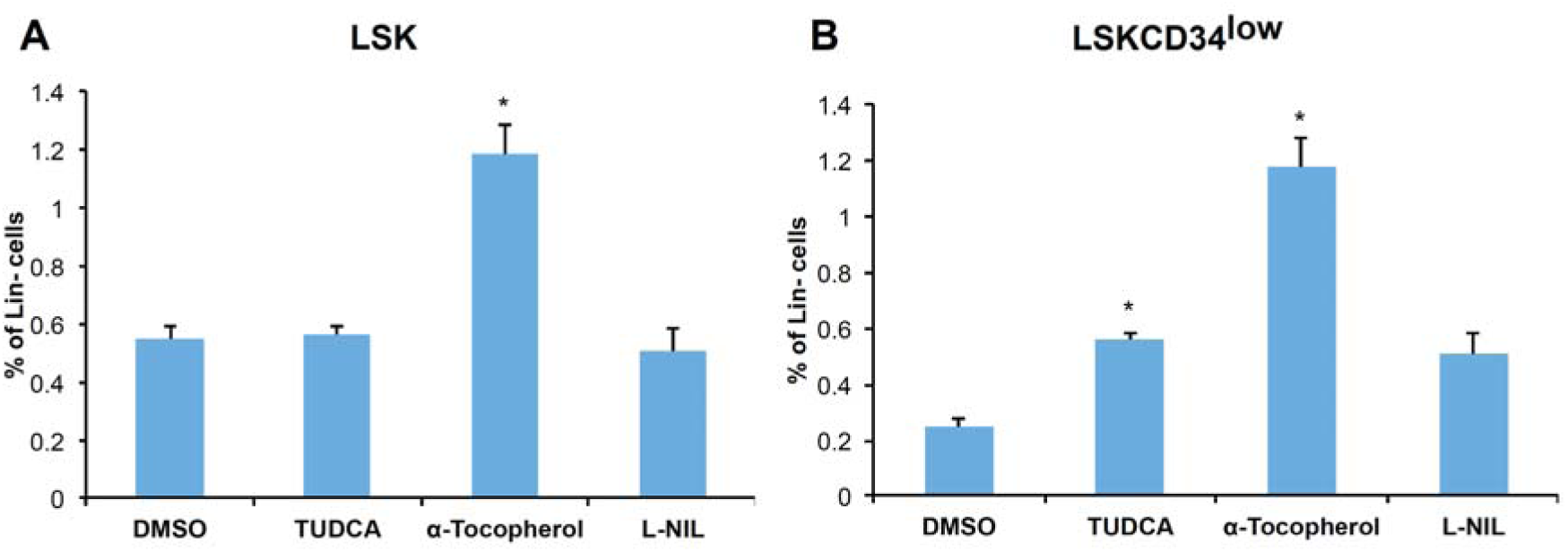
Effect of HSMs in HSC pool. Mouse lin-cells have been treated with DMSO (control, 0.5%), TUDCA (10 µM), α-Tocopherol (10 µM), L-NIL (10 µM) for 7 days and determined percent of C) Lin^-^Sca1^+^c-Kit^+^ (LSK), D) Lin^-^Sca1^+^c-Kit^+^CD34^low^ (LSKCD34^low^). * p < 0.05. n=3. (See related **Figure S8**)

Moreover, TUDCA, α-Tocopherol and L-NIL treatment of UCB mononuclear cells led to increased human HSPC content as measured by increased CD34+ and CD133+ cells (**Figure 9**). Next, we sought to determine the effect of HSMs in proliferation kinetics of BM-MSCs, AD-MSCs, HUVECs, and HDFs. We found that TUDCA, α-Tocopherol and L-NIL treatment lead to a significant increase in proliferation of HDFs (**Figure 10A**). However, we did not detect any significant change in proliferation of HUVECs or AD-MSCs compared to control with any of TUDCA, α-Tocopherol and L-NIL treatments (**Figure 10B and 10C**). Intriguingly, we found that L-NIL treatment decreased the proliferative rate of BM-MSCs (**Figure 10D**). Taken together, TUDCA, α-Tocopherol and L-NIL pose different properties in terms of cellular expansion that could be further exploited to determine a mixture of HSMs along with c-Myc inhibitor for robust and efficient HSPC expansion.

**Figure 9.**
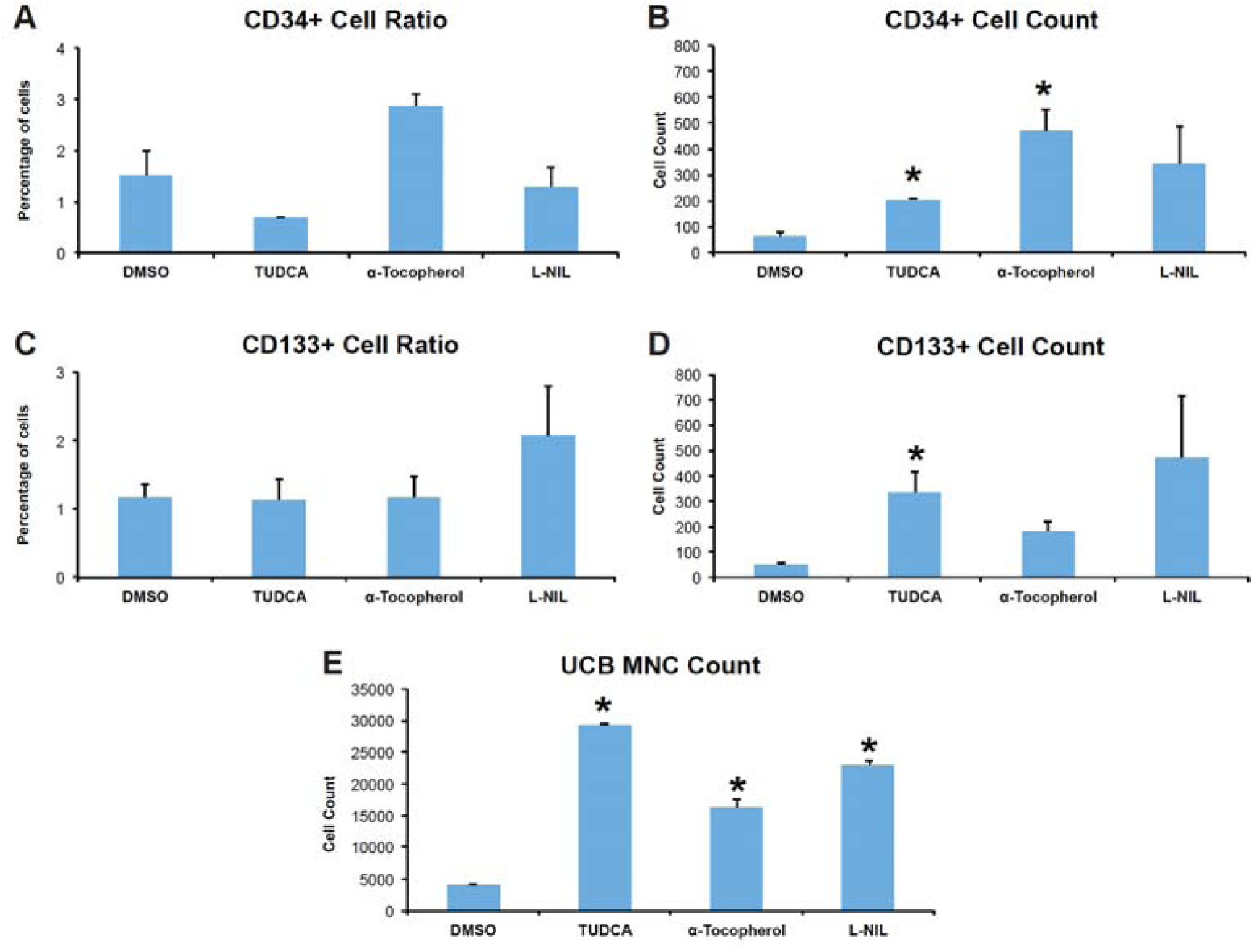
Effect of HSM treatments in human HSPC expansion. Human UCB MNCs have been treated with DMSO (control, 0.5%) TUDCA, α-Tocopherol, and L-NIL for 7 days and determined A) CD34+ cell ratio, B) CD34+ cell count, C) CD133+ cell ratio, C) CD133+ cell count, and E) UCB MNC count. * p < 0.05, n=3.

**Figure 10.**
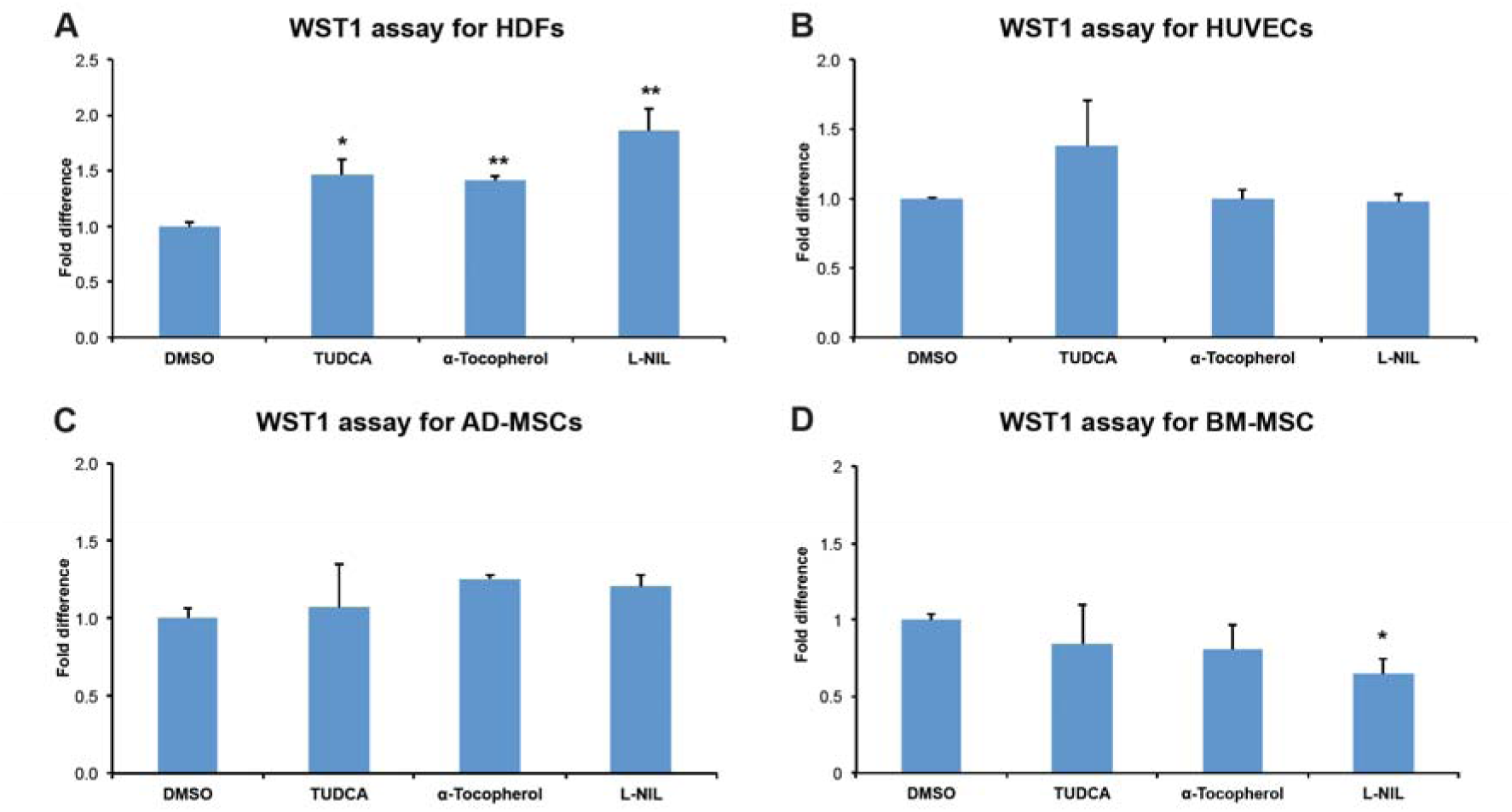
Effect of selected HSM treatments in endothelial cells, fibroblasts, AD-MSCs, and BM-MSCs. A) HDF, B) HUVECs, C) AD-MSC, and D) BM-MSC were treated with DMSO (control, 0.5%) and TUDCA, α-Tocopherol, L-NIL for 3 days. Cell proliferation was examined by WST1 assay.

### 3.8. Inhibition of c-Myc along with HSMs allows development of *ex vivo* HSC expansion cocktail with the concomitant improvement of HDR/S-phase gene expression

We hypothesized that an optimum combination of HSMs could provide a robust HSC expansion by targeting several pathways. In order to determine optimum combination of HSMs, we prepared combinatorial mixtures of TUDCA, α-Tocopherol, L-NIL along with c-Myc inhibitor 10074-G5 and tested their effect in HSC expansion (**Table 1**). The flow cytometry analysis of c-Kit+ **(Figure 11A)**, Sca1+ **(Figure 11B)**, LSK cells **(Figure 11C)**, and LSKCD34low HSCs **(Figure 11D)** demonstrated that c-Myc inhibitor 10074-G5 along with various HSMs (namely **M1**, **M3**, **M5**, and **M8** cocktails) led to improved maintenance and expansion of HSCs (**Figure 11A-D**). Combination of c-Myc inhibitor 10074-G5 along with TUDCA and L-NIL led to higher expansion of HSCs (**mixture M8**). This is evident in the induction of c-Kit+ alone, Sca-1+ alone, LSK, and LSKCD34^low^ cell content.

**Figure 11.**
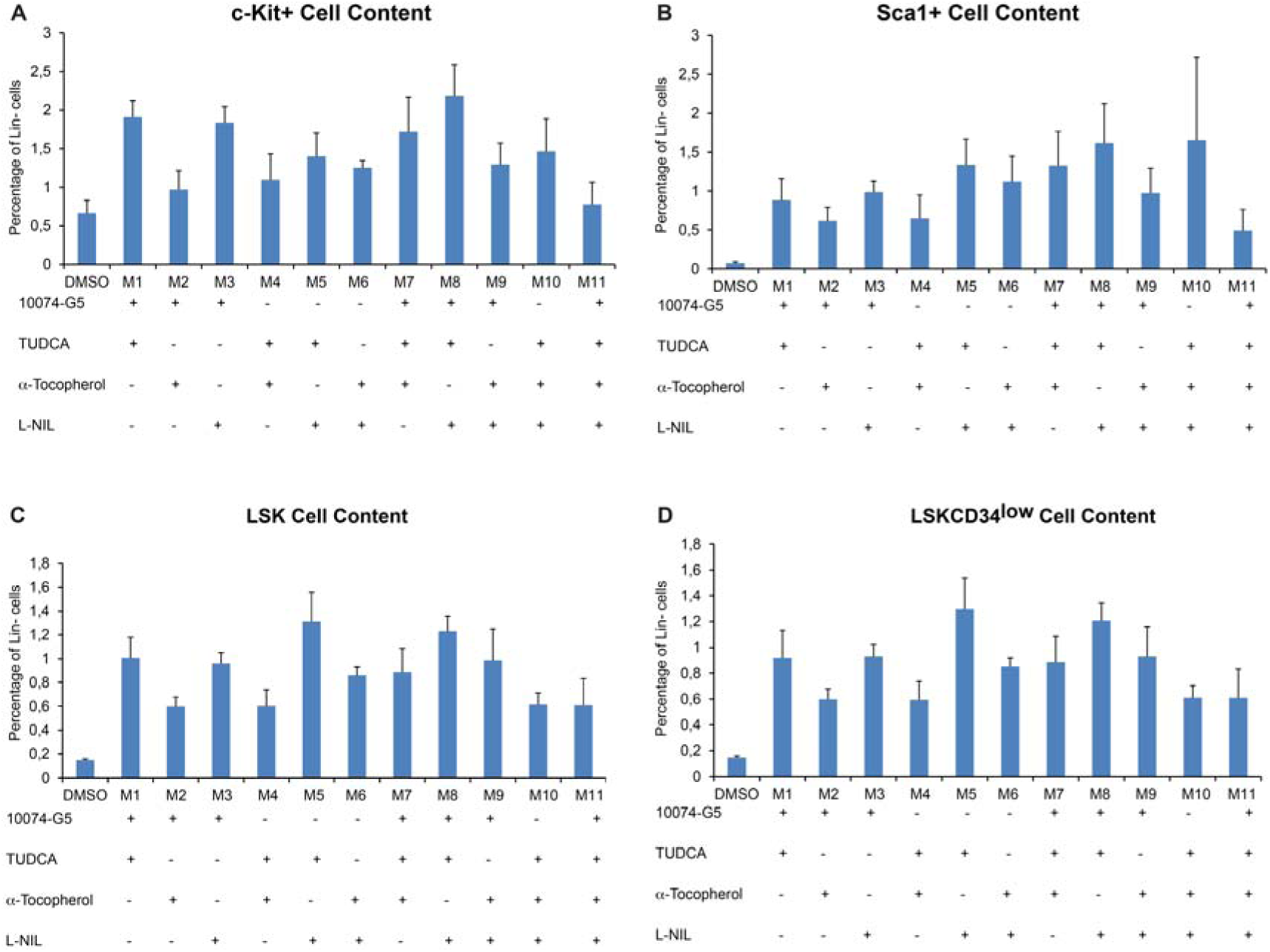
Combinatorial effect c-Myc inhibitor 10074-G5 with other HSMs. Mouse Lin-cells have been treated with DMSO (control, 0.5%) and combination of 10074-G5, TUDCA, α-Tocopherol, and L-NIL for 7 days. Ratio of A) c-Kit, B) Sca-1, C) Lin^-^Sca1^+^c-Kit^+^ (LSK), D) Lin^-^Sca1^+^c-Kit^+^CD34^low^ (LSKCD34^low^) cells have been determined. Note that **M8** treatment leads to induction of all HSC surface markers analyzed. * p < 0.05, ** p < 0.01, n=3.

Induction of homology directed repair (HDR) pathways and entry into S-phase of the cell cycle is crucial for efficient gene editing and proper expansion of HSCs. We show that **M8 cocktail** of 10074-G5, TUDCA, and L-NIL leads to improved HDR and S-phase related gene expression compared to DMSO and c-Myc inhibition alone (**Figure 12**). This is shown by induction of Rad51, Mre11a, and Mcm2 gene expression post treatments.

**Figure 12.**
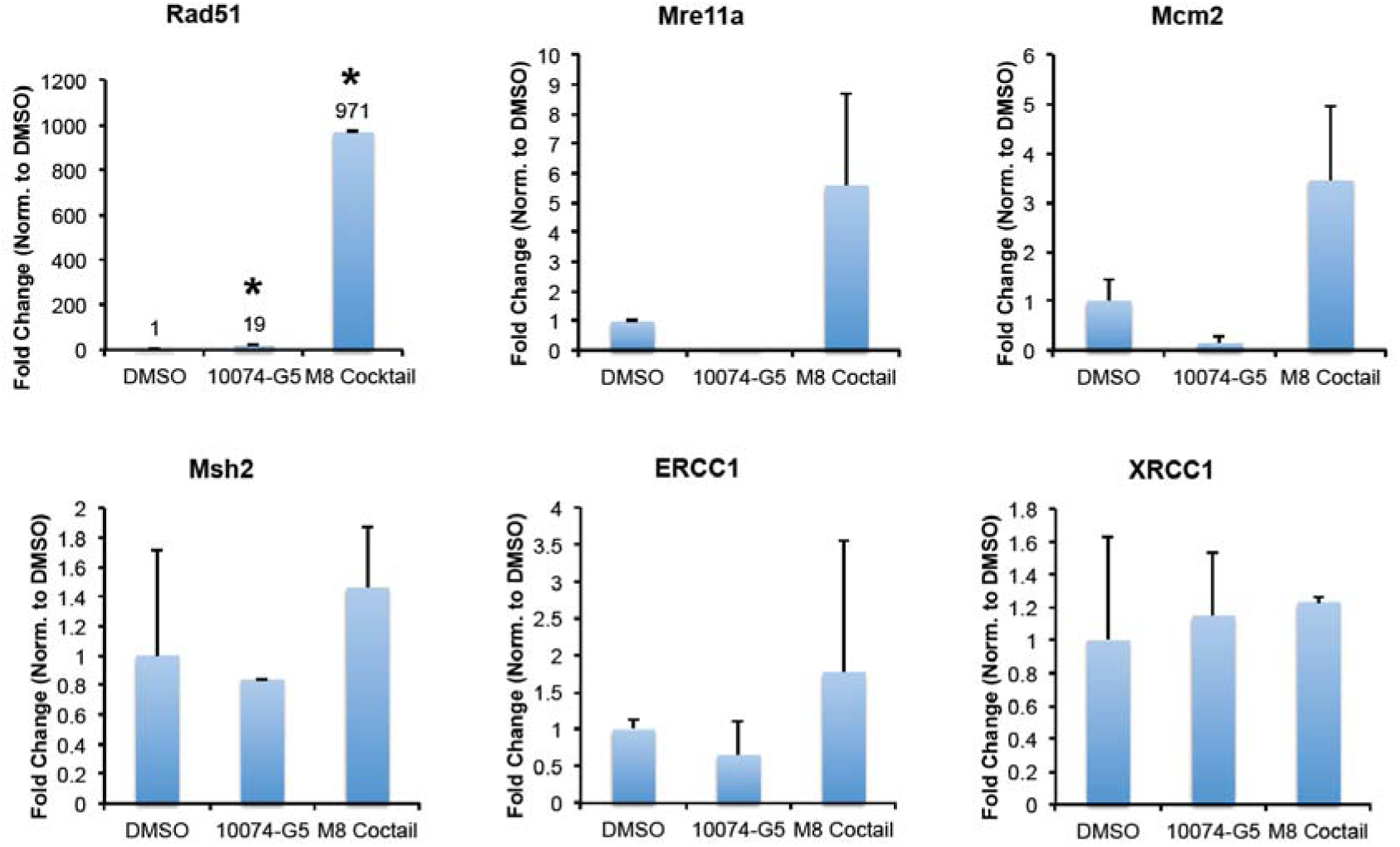
Analysis of HDR and S-phase gene expression post 10074-G5 and M8 coctail treatments. Mouse Lin-cells have been treated with DMSO (control, 0.5%), 10074-G5, and **M8 cocktail** of 10074-G5, TUDCA, and L-NIL for 7 days. Expression of HDR and S-phase progression related gene expression have been determined. Note that **M8** treatment leads to induction of key HDR and S-phase genes. * p < 0.05, n=3.

### 3.9. Repopulation of 10074-G5 and M8 treated HSCs in immunocomprimized animals

Engraftment and repopulation analysis have been carried out to determine HSC function in vivo. This is done to obtain in vivo information regarding functional ex vivo expansion of HSCs following treatment with DMSO, 10074-G5 or Mixture M8. For transplantation studies, we have used NOD/SCID mice homozygous for CD45.1 allele. Because donor cells used in ex vivo expansion procedure express CD45.2, engraftment and repopulation analysis of donor-derived blood cells studies have been accomplished by flow cytometric analysis of CD45.2+ cells in host animal. Here we show efficient bone marrow engraftment of 10074-G5 or Mixture M8 treated HSCs (**Figure 13A**). In addition, transplanted HSCs successfully repopulate whole blood lineages (B cells, T cells, Granulocytes, and Macrophages) (**Figure 13B**).

**Figure 13.**
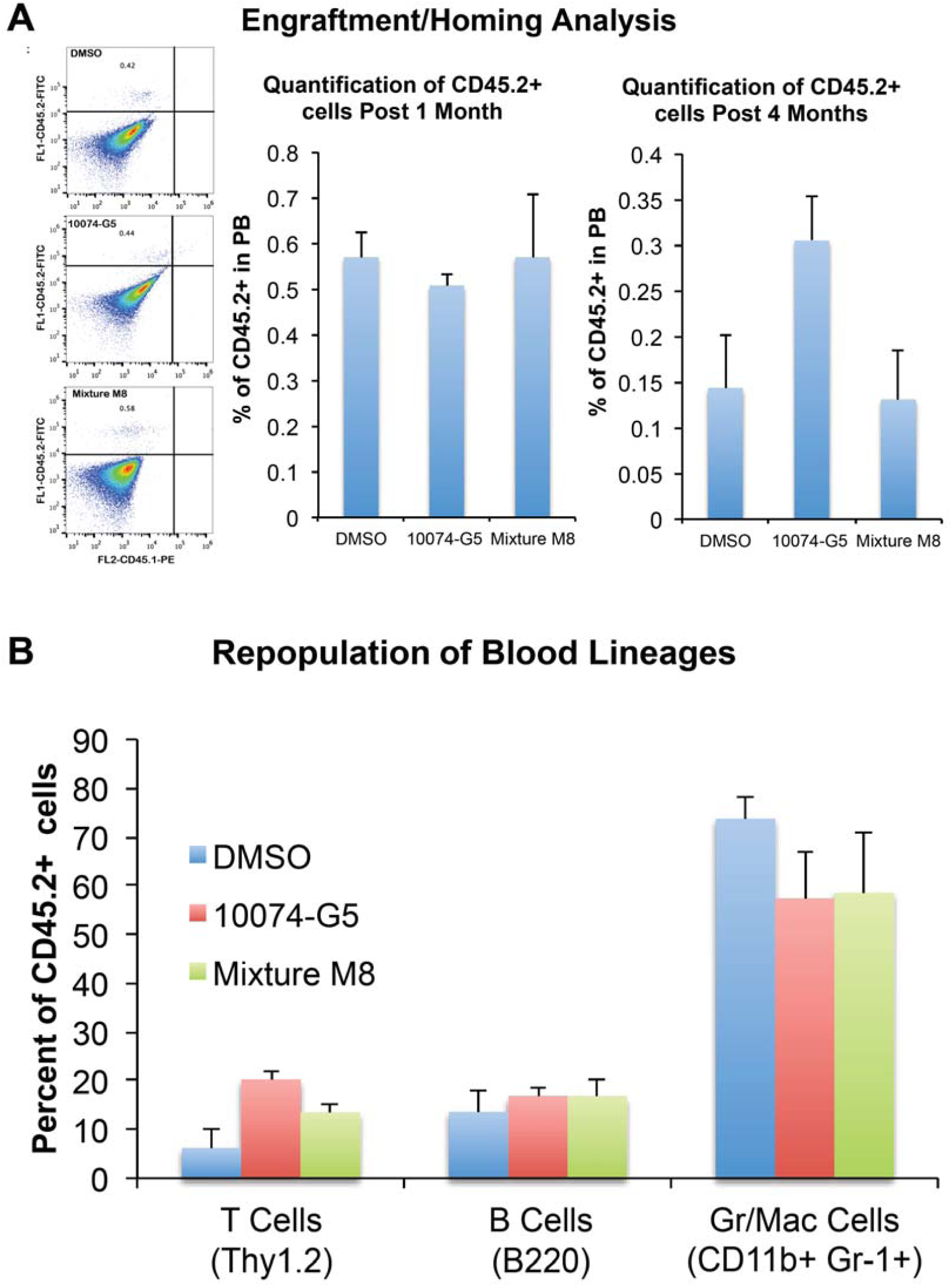
Analysis of engraftment and repopulation post 10074-G5 induced HSC transplantation. CD45.2+ HSCs are transplanted into CD45.1+ NOD/SCID animals and analyzed by flow cytometry for the engraftment and repopulation of blood linages post 1 and 4 months. **A)** Representative flow cytometry plots, and percent of CD45.2+ cells in the recipient mouse PB, **B)** Percentages of repopulating blood lineages. n=3.

## 4. DISCUSSION

Many studies addressed to define *ex vivo* culture conditions which relies on cytokines and growth factors to expand functional HSCs, however HSC expansion by utilizing small molecules with concomitant HDR modulation that targets HSC quiescence and metabolic regulators have not been widely studied. Previously, Wilson et al illustrated two to threefold rise in the Lin^-^ hematopoietic progenitors post genetic loss of c-Myc in a murine animal model (17). Thus, we targeted HSC regulator c-Myc using small molecule inhibitor 1007-G5. 1007-G5 is a specific and potent inhibitor of c-Myc specific DNA interaction and target gene regulation. Intriguingly, c-Myc inhibitor 10074-G5, which has not been previously studied in small molecule induced *ex vivo* murine and human HSC expansion procedure, significantly increased murine LSK and LSKCD34^low^ HSC, and human CD34+ and CD133+ HSC compartments.

In the present study, it has been demonstrated that c-Myc inhibitor promoted proliferation of normally quiescent hematopoietic stem and progenitor cells. This is further supported by lower expression of CDKIs both *ex vivo* and *in vivo* post c-Myc inhibition. In addition, c-Myc modulates glycolysis through transcriptional regulation of key genes involved in. c-Myc activates the expression of lactate dehydrogenase A (LDHA), which generates lactate from pyruvate as part of the glycolytic pathway. In addition, several glucose metabolism genes are directly regulated by c-Myc, including glucose transporter (GLUT1), phosphofructokinase (PFKM), hexokinase 2 (HK2), and enolase 1 (ENO1) (28-31). Through the upregulation of these downstream targets, c-Myc induces the Warburg effect, which enables generation of pyruvate from glucose even under low oxygen tension. In this study, we demonstrated that reduced glycolytic gene expression may shed light to increased expansion of HSCs. Previously we and other studies showed that HSCs reside in hypoxic niches with preferential utilization of glycolysis through involvement of Hif-1α and Hif-2α under transcriptional regulation of Meis1 (21). We have shown that HSC was expanded *in vivo* after stem cell specific knockout of hematopoietic factors Meis1 and Hif-1α (7). Deletion of such HSC hypoxic niche related factors in HSC compartment not only lead to altering preference away from glycolysis to mitochondrial phosphorylation in HSCs, but also cell cycle entry and expansion of HSC pool. Similarly, c-Myc inhibition might modulate HSC metabolism away from niche preferred cytoplasmic glycolysis, thus allowing exit from metabolic quiescent state and then get metabolically activated. However, further studies needed to determine how c-Myc inhibition alters metabolic preference of HSCs.

Two stem cell populations located in adult bone marrow are hematopoietic stem cells and mesenchymal stem cells. Previous studies showed that c-Myc has been shown to play a role in HSCs to exit from the stem cell niche through the down-regulation of N-cadherin (17, 32). In this study by targeting c-Myc with small molecules, however, we could not detect any change in the extent of HSC mobilization from bone marrow to peripheral blood. Intriguingly, while c-Myc inhibitor 10074-G5 increased *in vivo* murine HSC content in the bone marrow, it inhibited the proliferation of BM-MSCs. On the other hand, there were no effect on the proliferation kinetics of AD-MSCs, endothelial cells or fibroblasts. BM-MSCs have ability differentiate osteoblasts, which are important modulators of HSC niche (33). These findings suggest that *in vivo* osteoblastic bone marrow mesenchymal and stem cell niche could be altered following c-Myc inhibition.

Long-term HSC expansion procedures involving growth factors and cytokines often result in HSC exhaustion. This is could be overcome with approaches that target factors leading to senescence and cell cycle exit as shown with inhibition of p38 (34). Moreover, cytokines including thrombopoietin (TPO), FL3, IL3, IL6, IL11 and stem cell factor (SCF) have been proved to have function in HSC expansion (2). They stimulate HSCs that are arrested in G_0_ phase to enter the cell cycle by up-regulating factors responsible in self-renewal and by down-regulating inhibitors of cell cycle. Small molecules have assisted the exploration of the signaling pathways which regulate stemness and been used to expand HSCs *ex vivo* (5). In addition, cointroduction of hematopoietic small molecules could provide a mean to expand HSCs by targeting several distinct pathways. We show that c-Myc inhibition alters expression of glycolysis-associated genes. TUDCA treatment enhances protein folding, resulting in reduced ER stress (35-37). L-NIL, on the other hand, is selective iNOS inhibitor and associated with lower apoptosis in CD34+ HSCs (38). Thus, we have shown that combination of c-Myc inhibition along with TUDCA and L-NIL treatment provided the robust HSC expansion through modulation of different mechanisms.

We have also explored the extent of modulation of homology directed repair (HDR) pathways and entry into S-phase of the cell cycle, which are crucial for efficient gene editing and proper expansion of HSCs. Cocktail of 10074-G5, TUDCA, and L-NIL lead to enhanced HDR and S-phase related gene expression in hematopoietic cells. We have found Rad51, which has central role in homologous recombinational repair and DNA double strand break repair, with most robust induction compared to DMSO or c-Myc inhibition alone. RAD51 recombinase modulates strand transfer between a DNA double strand break sequence and its undamaged homologue to allow repair of the mutated or damaged region through homology based re-synthesis (39, 40). In addition, improved HDR and S-phase progression supported by induction of double-strand break repair protein Mre11a and Mcm2 gene expression. Mre11a is plays important role in involved in homologous recombination, telomere length maintenance, and DNA double-strand break repair(41, 42). Mcm2, mini-chromosome maintenance protein 2, is known as DNA replication licensing factor (43, 44). Mcm2 is key component of pre-replication complex in the S-Phase and modulate helicase activity of MCM complex(45, 46). Upregulation of HDR and S-Phase related gene expression suggest that cocktail of 10074-G5, TUDCA, and L-NIL could increase efficiency of CRISPR/CAS9 mediated gene editing through activation of HDR (47-50). This cocktail of HSM warrants further studies in HSC expansion post gene editing in BM derived HSCs and expansion of sufficient number of cord blood HSCs for allogeneic HSC transplantation into adult recipients.

## Acknowledgments

We thank the support from the European Commission Co-Funded Brain Circulation Scheme by The Marie Curie Action COFUND of the 7th. Framework Programme (FP7) and The Scientific and Technological Research Council of Turkey (TÜBİTAK) [grant number 115C039], TÜBİTAK ARDEB 1001 [grant numbers 115S185 & 215Z069], TÜBİTAK ARDEB 3501 [grant number 215Z071], The Science Academy Young Scientist Award Program (BAGEP-2015, Turkey), The International Centre for Genetic Engineering and Biotechnology – ICGEB 2015 Early Career Return Grant [grant number CRP/TUR15-02_EC], Medicine for Malaria Venture MMV Pathogenbox Award (Bill and Melinda Gates Foundation) and funds provided by Yeditepe University, Istanbul, Turkey. EA has been supported by TÜBITAK-BIDEB 2211 program. MA has been supported by TÜBITAK-BIDEB 2213 program.

## Author Contributions

MA, EA, DDC, GSA, RDT, LYA, PS, ECT, SC, DY, NM, ZG, FS, FK designed studies, performed experiments and prepared figures. MA and FK wrote the main manuscript text. All authors reviewed the manuscript.

## Figures and Figure Legends

**Figure S1.**
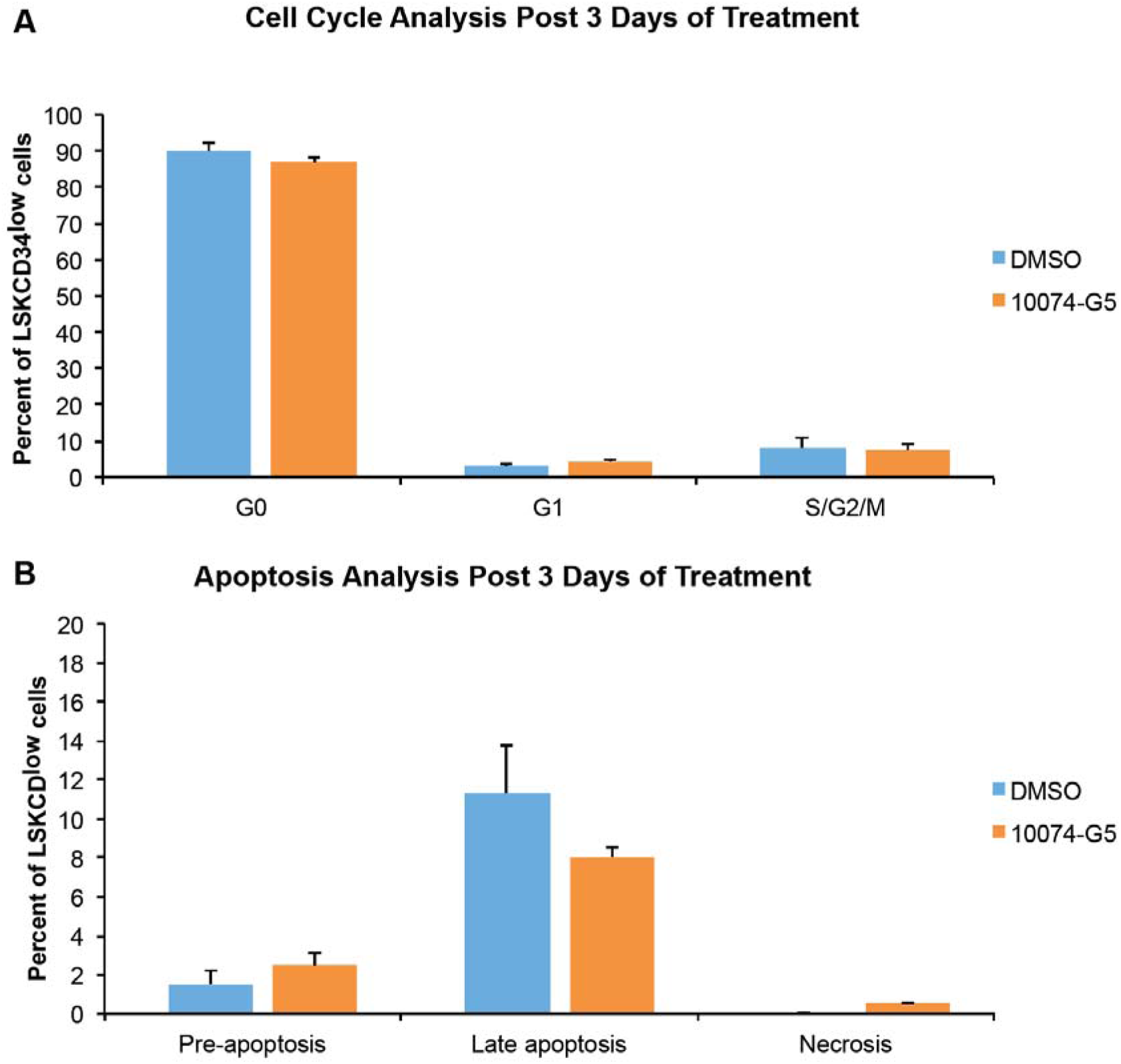
Murine HSCs have been treated with DMSO (control, 0.5%), c-Myc inhibitor 10074-G5 (10 µM) for 3 days. Analysis of **A)** cell cycle and **B)** apoptosis. n=3. (See related **Figure S9**)

**Figure S2.**
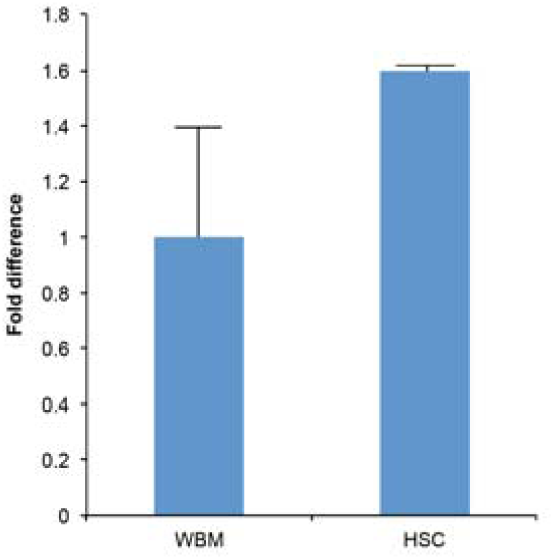
Expression profile of c-Myc in whole bone marrow (WBM) cells versus HSCs. n=2.

**Figure S3.**
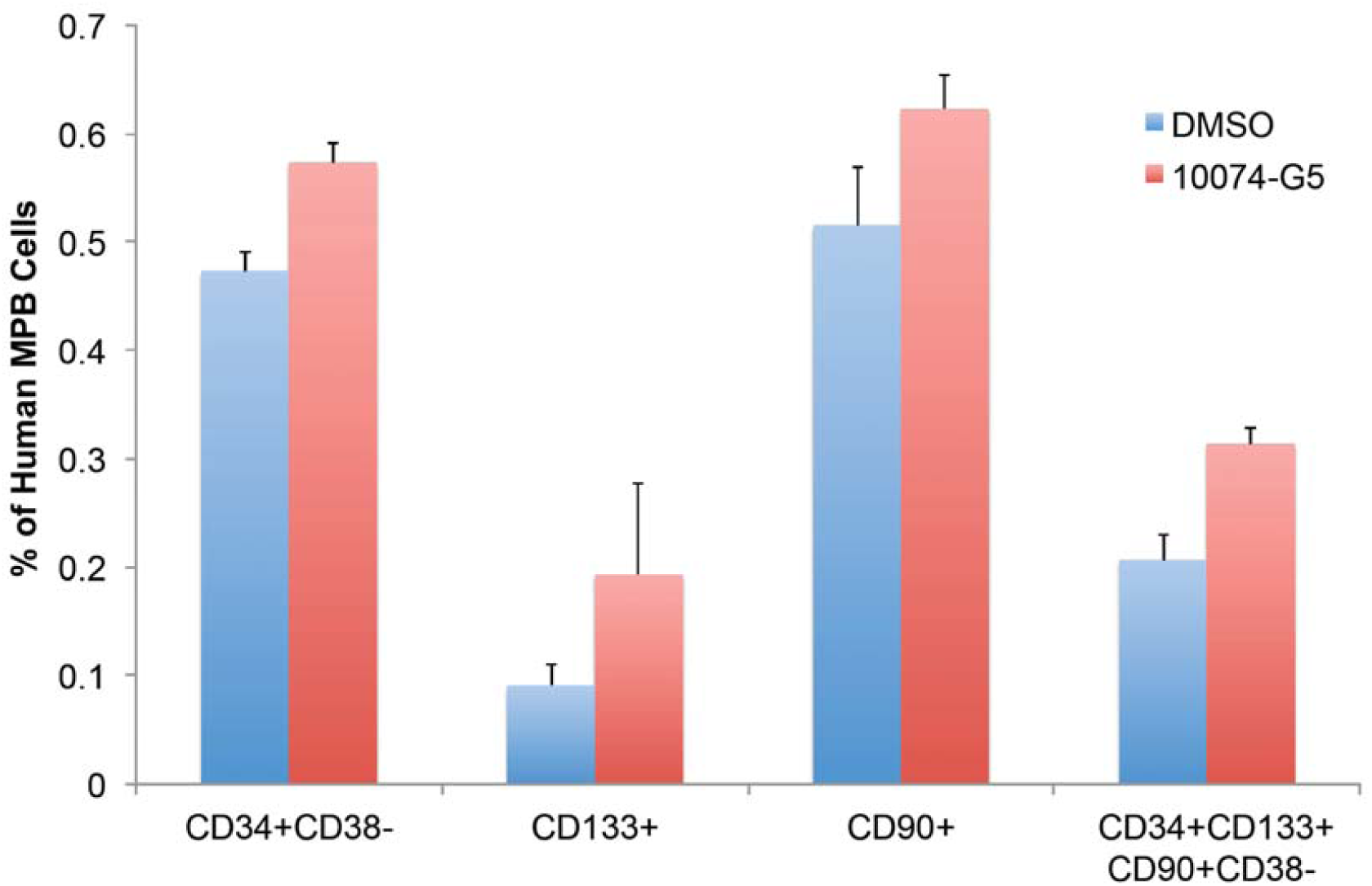
Analysis of human HSC content in mobilized peripheral blood (MPB) cells post c-Myc inhibition. n=3.

**Figure S4.**
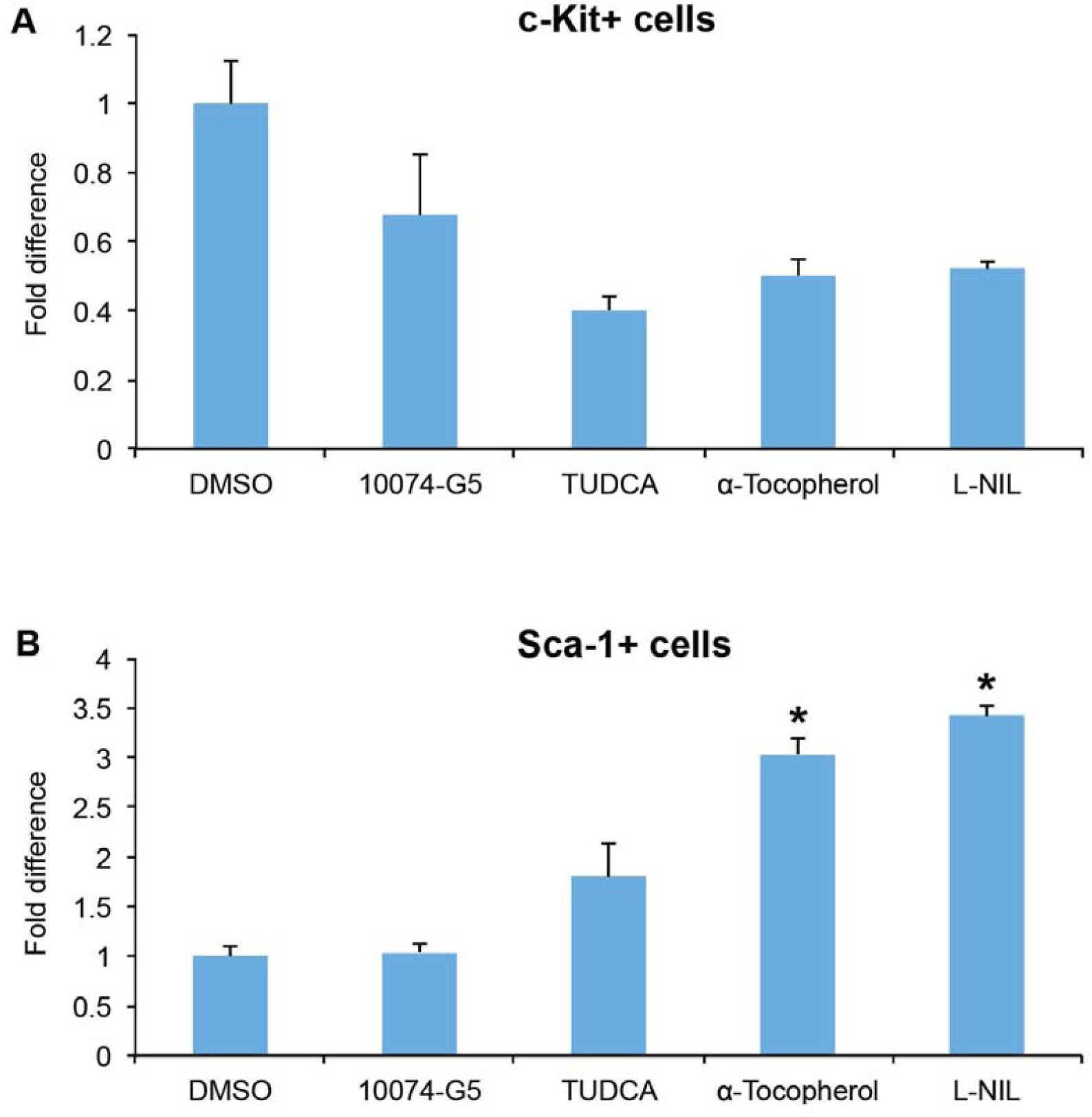
Effect of HSM treatments in c-Kit+ and Sca1+ cell content. Murine lin-cells have been treated with DMSO (control, 0.5%), c-Myc inhibitor 10074-G5, TUDCA (10 µM), α-Tocopherol (10 µM), L-NIL (10 µM) for 7 days and determined percent of **A)** c-Kit+ and **B)** Sca-1+ cells. * p < 0.05, n=3.

**Figure S5.**
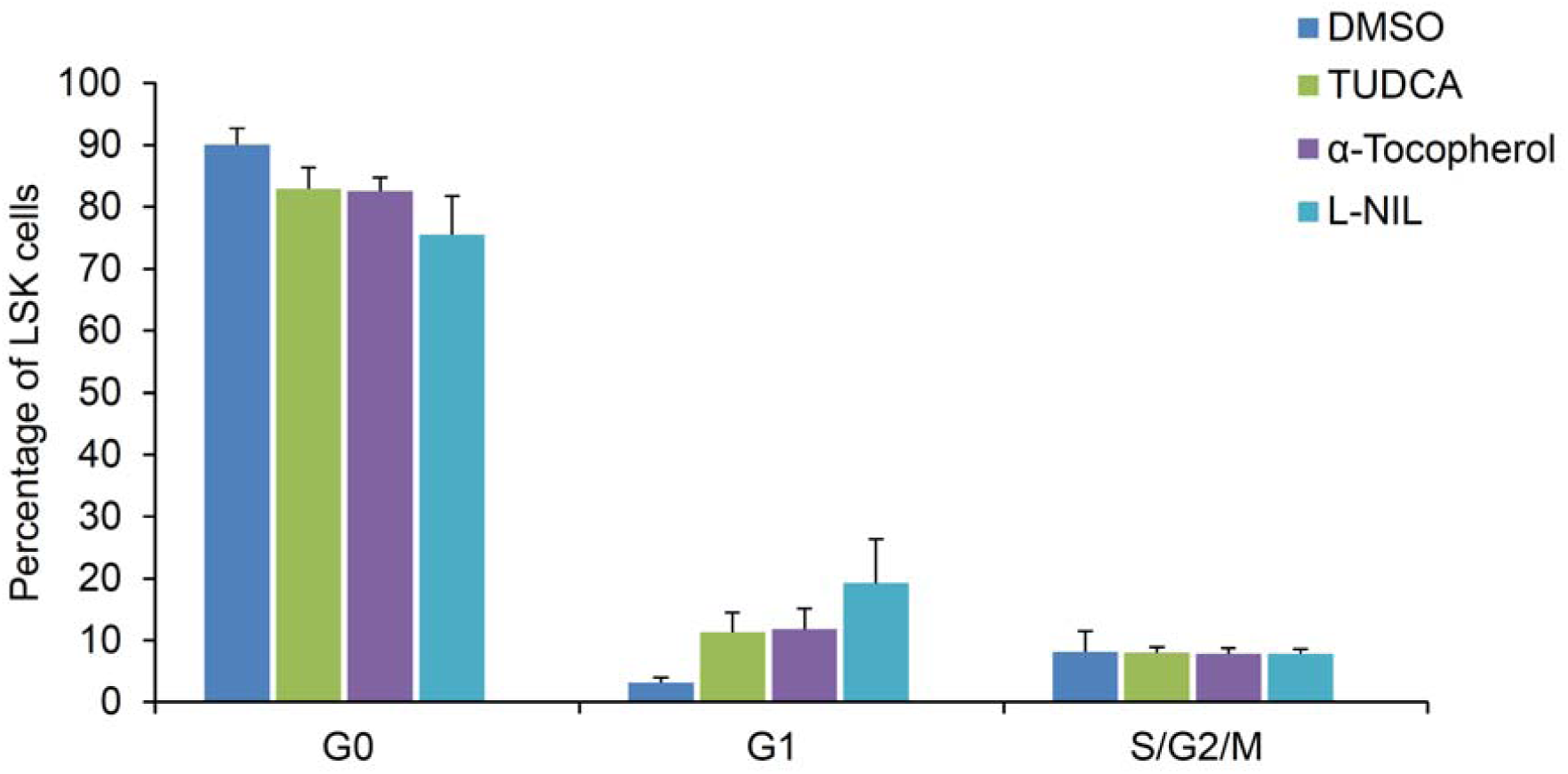
Cell cycle analysis of HSCs after HSM treatments. Murine HSCs have been treated with DMSO (0.5%), TUDCA (10 µM), α-Tocopherol (10 µM), L-NIL (10 µM) for 3 days and determined percent of cells in G_0_, G_1,_ and S/G_2_/M. n=3.

**Figure S6.**
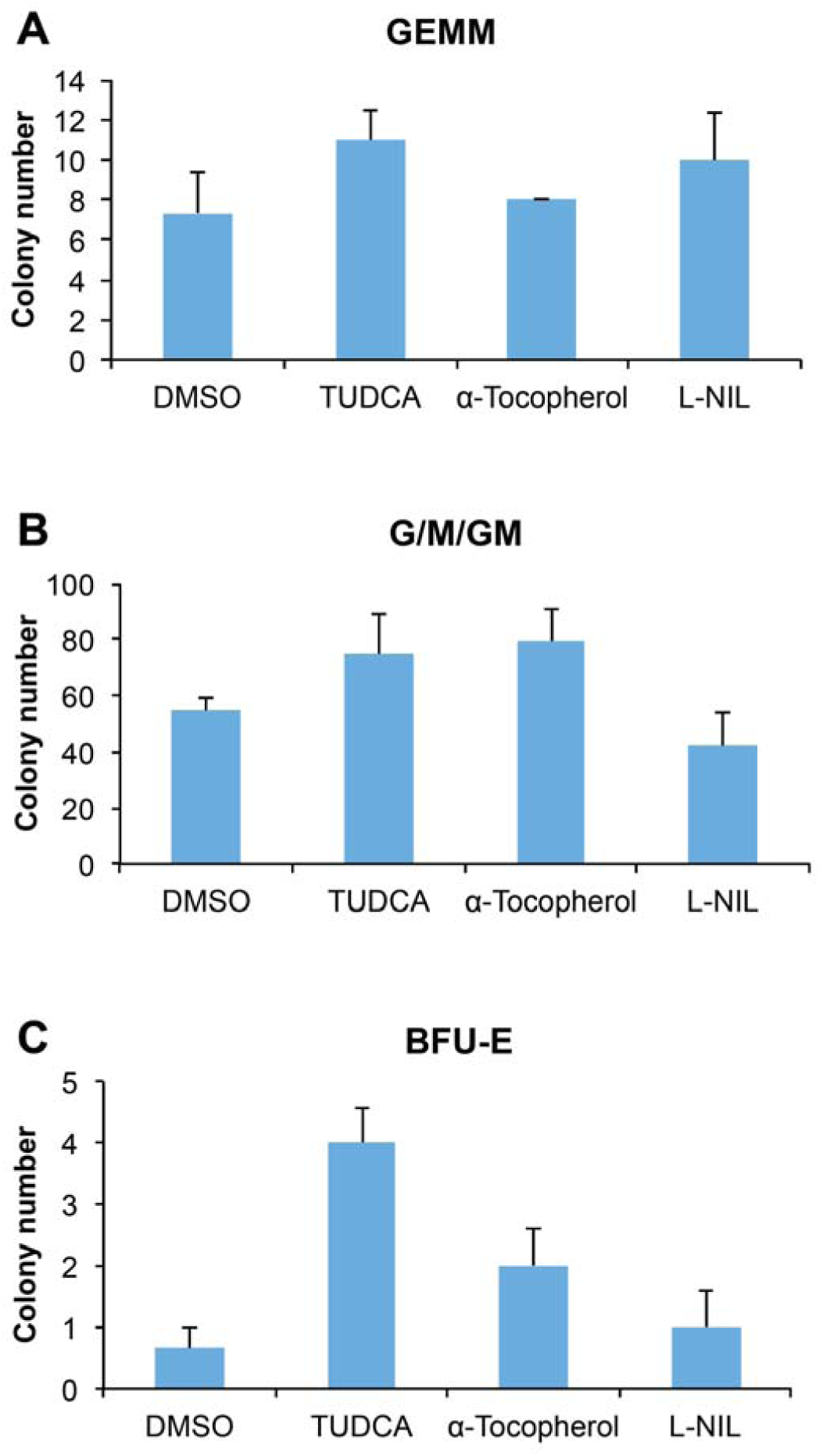
Apoptosis analysis of murine HSCs after HSM treatments. Murine HSCs have been treated with DMSO (control, 0.5%), TUDCA (10 µM), α-Tocopherol (10 µM), L-NIL (10 µM) for 3 days followed by staining with Annexin V FITC and propidium iodide and determined percent of pre-apoptotic, late apoptotic and necrotic cells. n=3.

**Figure S7.**
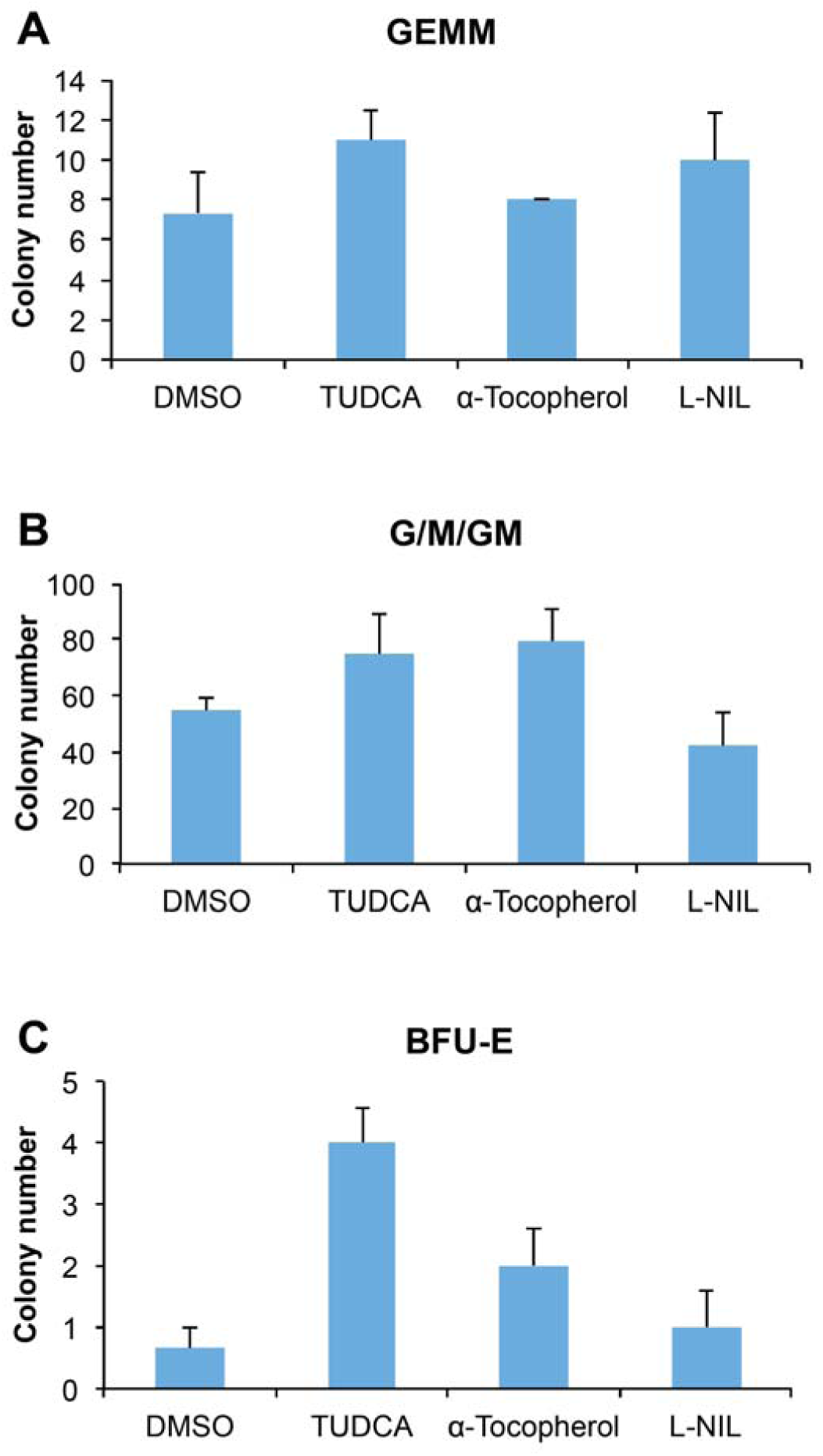
CFU analysis of HSCs post HSM treatments. Number of A) GEMM, B) G/M/GM, C) BFU-E colonies formed post 13 days. n=3.

**Figure S8.**
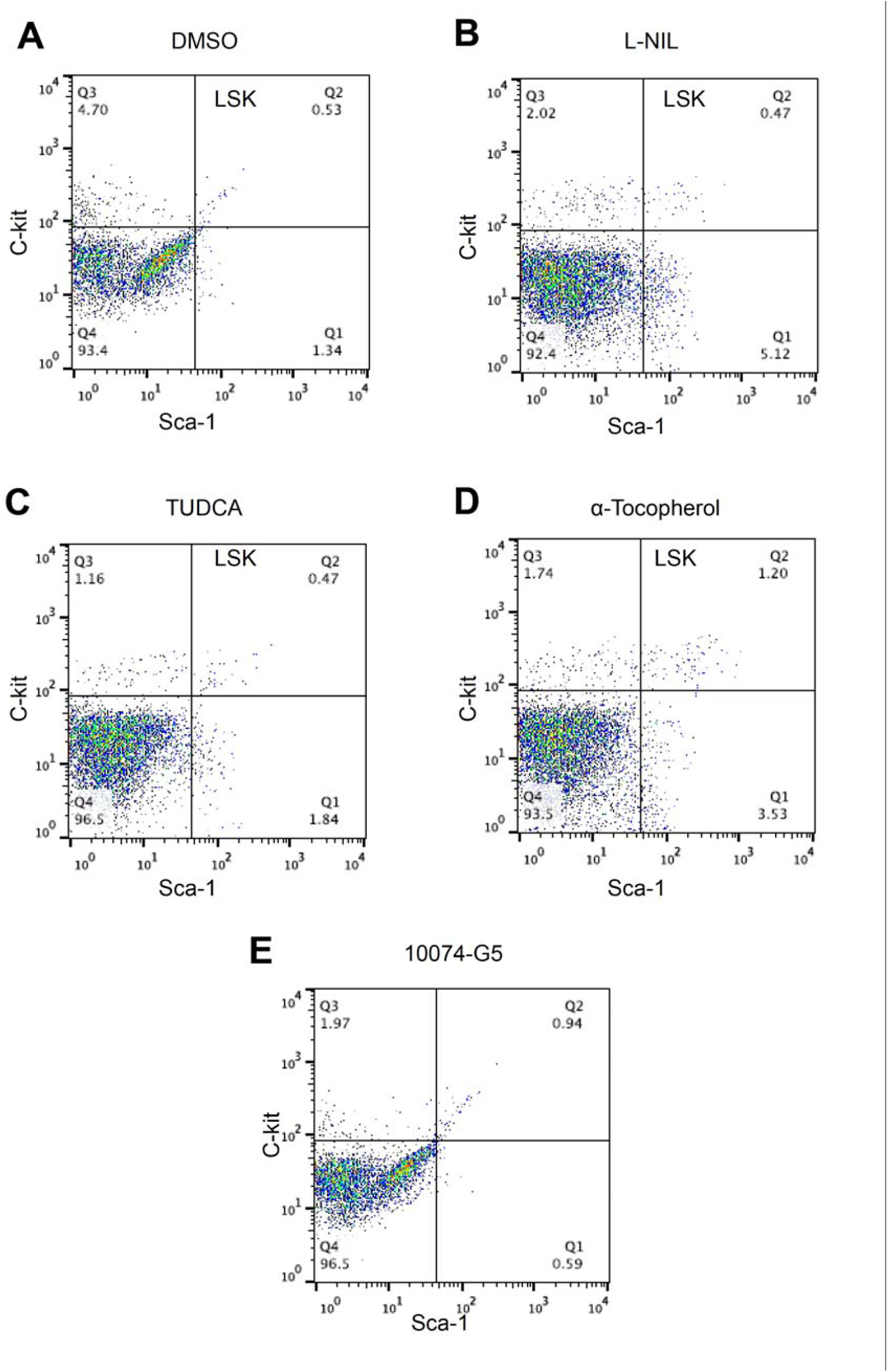
Flow cytometry plots of small molecule treated cells

**Figure S9.**
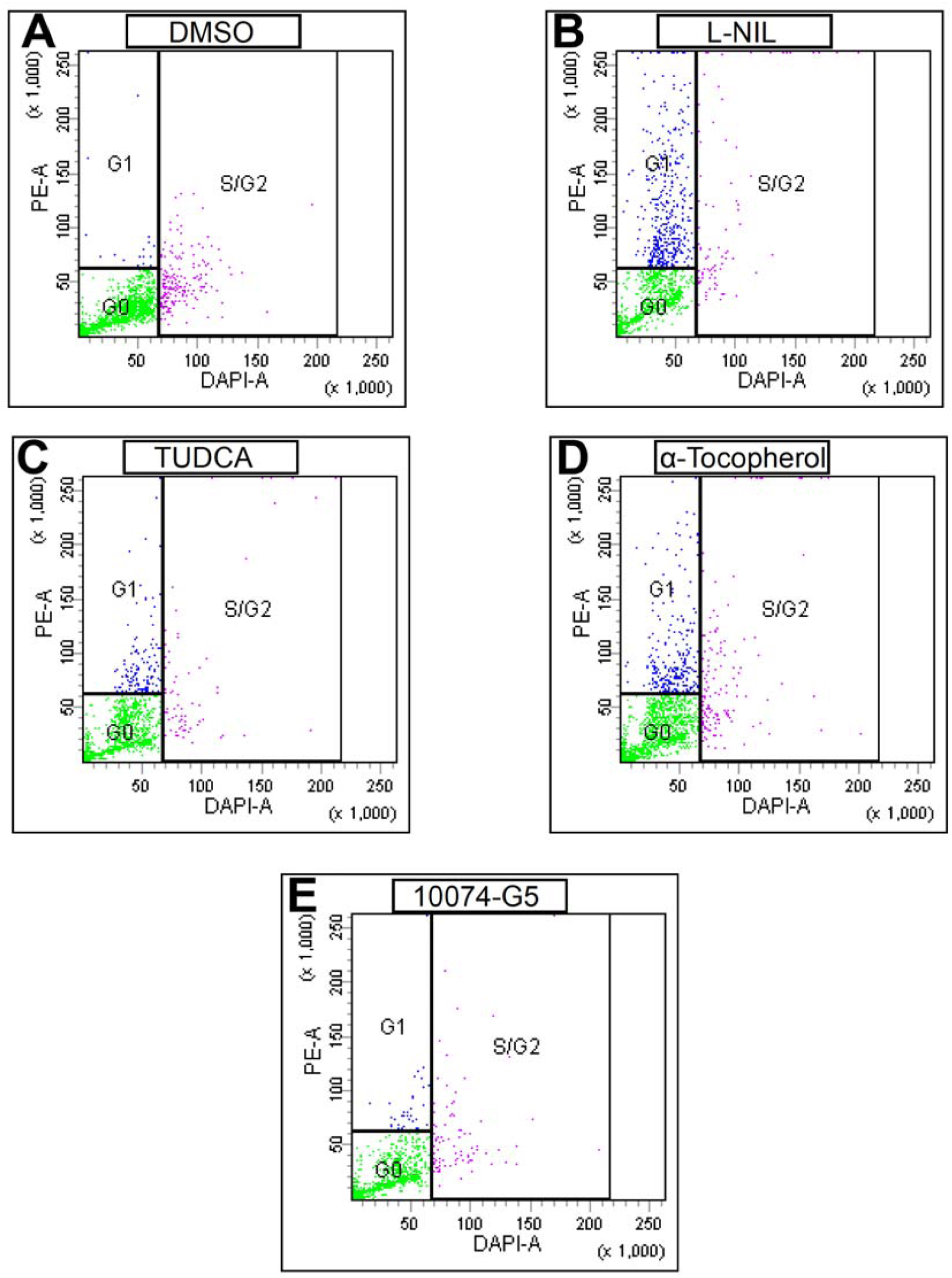
Cell Cycle analysis by Flow cytometry. Representative plots of small molecule treated cells

